# The genetic status of IDH1/2 and EGFR dictates the vascular landscape and the progression of gliomas

**DOI:** 10.1101/2020.09.21.306134

**Authors:** Berta Segura-Collar, María Garranzo-Asensio, Beatriz Herranz, Esther Hernández-SanMiguel, Bárbara. S. Casas, Ander Matheu, Ángel Pérez-Núñez, Juan M. Sepúlveda-Sánchez, Aurelio Hernández-Laín, Verónica Palma, Ricardo Gargini, Pilar Sánchez-Gómez

## Abstract

**Rationale:** Glioma progression is driven by the induction of vascular alterations but how the tumor genotype influence these changes is still a pending issue. We propose to study the underlying mechanisms by which the genetic changes in *isocitrate dehydrogenase 1/2* (*IDH1/2*) and *epidermal growth factor receptor* (*EGFR*) genes establish the different vascular profiles of gliomas.

**Methods:** We stratified gliomas based on the genetic status of *IDH1/2* and *EGFR* genes. For that we used in silico data and a cohort of 93 glioma patients, where we analyzed the expression of several transcripts and proteins. For the in vitro and in vivo studies, we used a battery of primary glioblastoma cells derived from patients, as well as novel murine glioma cell lines expressing wild-type or mutant EGFR. In these models, the effect of the small molecule ibrutinib or the downregulation of CD248 and SOX9 was evaluated to establish a molecular mechanism.

**Results:** We show that *IDH1/2* mutations associate with a normalized vasculature. By contrast, *EGFR* mutations stimulate the plasticity of glioma cells and their capacity to function as pericytes in a bone-marrow and X-linked (BMX)/SOX9 dependent manner. The presence of tumor-derived pericytes stabilize the profuse vasculature and confers a growth advantage to these tumors, although they render them sensitive to pericyte-targeted molecules. Wild-type/amplified *EGFR* gliomas are enriched in blood vessels too, but they show a highly disrupted blood-brain-barrier due to a decreased BMX/SOX9 activation and pericyte coverage. This leads to poor nutrient supply, necrosis and hypoxia.

**Conclusions:** The function of tumor-derived pericytes delimitates two distinct and aggressive vascular phenotypes in *IDH1/2* wild-type gliomas. Our results lay the foundations for a vascular dependent stratification of gliomas and suggest different therapeutic vulnerabilities depending on the genetic status of *EGFR*.

Graphical Abstract. Schematic view of IDH and EGFR function in the regulation of glioma microenvironment.
Mutant IDH gliomas express low levels of angiogenic molecules and have a vasculature reminiscent of normal tissue. EGFR mutations drive glioma growth by promoting tumor-to-pericyte transdifferentiation and vascular stabilization in a BMX-SOX9 dependent way. Leaky vessels with hypoxia and necrosis characterize tumors overexpressing the wild-type isoform of the receptor. These phenotypes determine the response to therapy of the different IDH wild-type gliomas.

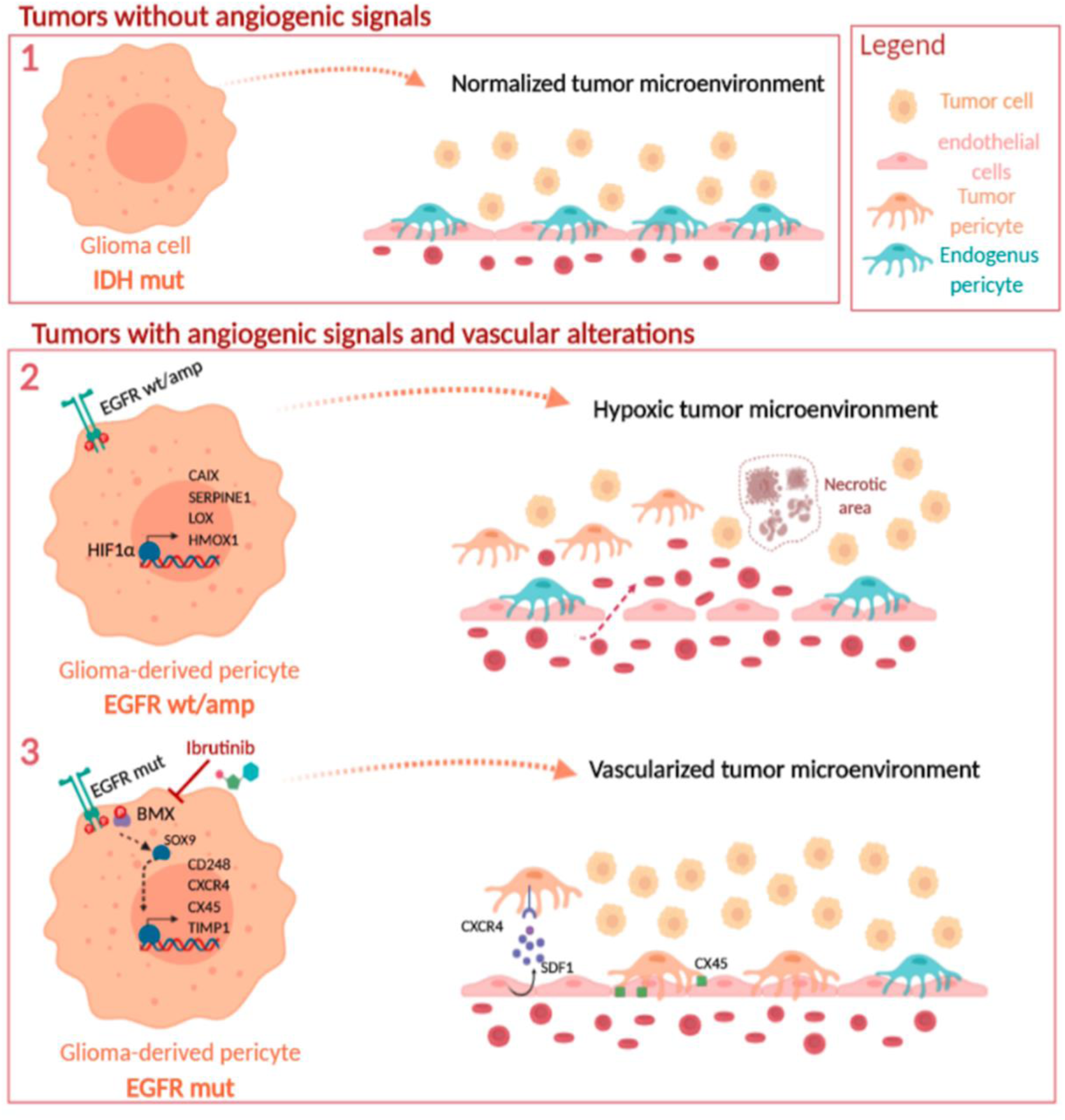

## Introduction

Diffuse gliomas are histologically classified as low and intermediate-grade gliomas (Lower-Grade Gliomas, LGG) (grades 2 and 3) or glioblastomas (GBMs) (grade 4) (1). The poor prognosis of these tumors has been attributed in part to treatment limitations related to the tumor localization, but also to the insufficient knowledge about their physiology.

Florid vascular proliferation and aberrant vasculature are distinctive pathological hallmarks of glioma progression. Four distinct mechanisms of vascularization have been proposed in gliomas: vascular co-option, angiogenesis, vasculogenesis and vascular mimicry. The last one is associated with the plasticity of GBM cells, which can transdifferentiate into endothelial cells (ECs) (2) and pericytes (3). All these mechanisms are inter-linked and overlapped in the history of gliomas and give rise to an abnormal vascular network, with dilated and tortuous blood vessels (BVs) and poor pericyte coverage, which contributes to the formation of an abnormal blood-brain-barrier (BBB) (4).

Whereas the mechanisms that drive glioma neo-vascularization have been described, the genetic alterations that govern them have not been well established. However, it is known that mutations in the isocitrate dehydrogenase 1 and 2 (*IDH1/2*) genes, commonly found in LGG, favor the normalization of the vasculature. IDH mutant (IDHmut) gliomas have smaller BVs and are associated with less hypoxia compared with their wild-type (wt) counterparts (5). Moreover, we have recently described that the microtubule stabilizer TAU/MAPT is induced in IDHmut gliomas and promotes vascular normalization by opposing EGFR signaling (6).

The *EGFR* gene is amplified in 50 to 60% of IDHwt GBMs and half of these tumors carry the vIII variant (exons 2-7 deletion), which generates a constitutive activation of the receptor’s tyrosine kinase. In addition, missense point mutations in the extracellular domain of the receptor are also commonly found in gliomas (7). All these mutations confer a higher oncogenic potential to the tumor cells through the increase in their proliferative capacity. Moreover, EGFR activation in glioma cells promotes STAT3/5 activation and cytokine secretion to modify the microenvironment and thus drives tumorigenesis (8). This adds to the idea that the genotype of glioma driving cells determines their surrounding stroma. Here, we investigate how alterations in *EGFR* affect the vascular phenotype of gliomas and the implications for the progression of the tumors and for their susceptibility to therapy.

## Materials and Methods

### Human samples

The primary cell lines (Table S1) belong to the Biobank of that Hospital. Fresh tissue samples were digested enzymatically using Accumax (Millipore) and were maintained in stem cell medium; Neurobasal (Invitrogen) supplemented with B27 (1:50) (Invitrogen); GlutaMAX (1:100) (Invitrogen); penicillin-streptomycin (1:100) (Lonza); 0.4% heparin (Sigma-Aldrich); and 40 ng/ml EGF and 20 ng/ml bFGF2 (Peprotech).

### Mouse cell lines

Mouse SVZ cell lines were obtained by retroviral expression of EGFRwt or EGFRvIII in primary neural stem cell (NSC) cultures from the Subventricular Zone (SVZ) of p16/p19 ko mice. The NSCs were obtained as previously described (9) and they were grown in stem cell medium. After infection, the cells were injected into C57/BL6 mice. The tumors that grew were dissociated and the lines SVZ-EGFRwt/amp and SVZ-EGFRvIII were established. Both models express GFP and luciferase as a reporter. The GL261 murine glioma cells were maintained in DMEM plus 10% FBS, 2mM L-glutamine, 0.1% penicillin (100 U/ml) and streptomycin (100 μg/ml).

### DNA constructs and lentiviral/retroviral production

Retroviral vectors used were pBabe-EGFR wt (#11011) and MSCV-XZ066-GFP-EGFR vIII (#20737). pLV-Hygro-Luciferase (VectorBuilder #VB150916-10098). Lentiviral vector to express shRNAs were: shCD248 (Sigma #SHCLNG-NM_020404: TRCN0000053455, TRCN0000053457, TRCN00000443679, TRCN00000429396, TRCN0000043782) and shSOX9 (Adgene #40644).

### In vitro treatments

Cells were treated with ibrutinib (MedChemExpress, 31976) 5 µM ibrutinib, DMSO (control), MG132 (Millipore) 10 µM, or different concentrations of dacomitinib (Pfizer, PF-299804) or vehicle (DMSO).

### In vivo assays

Intracranial or subcutaneoous transplantations were establish as previously described (10). Mice were treated with ibrutinib at 12 mg/kg/day through intraperitoneal injection (i.p.), or dacomitinib (Pfizer, PF-299804) at 15mg/Kg/day (i.p.). For drug preparation ibrutinib was dissolved in 4% DMSO + 10% Hydroxypropyl-β-Cyclodextrin (HP-β-CD) and dacomitinib was dissolved in 20mM sodium lactate (pH=4) (1,5mg/ml). Control animals were treated with these solvents.

### Mouse magnetic resonance imaging (MRI)

Before imaging, animals were anesthetized using 2 % isofluorane (Isobavet, Schering-Plough) and they were IP injected with 0,1 mL of Gd-DOTA (Dotarem, Guerbet). Images were acquired on a 4,7 T Biospec BMT 47/40 spectrometer (Bruker), equipped with a 6 cm actively shielded gradient system, capable of 450 mT/m gradient strength (Universidad Complutense CAI facility).

### Cell sorting

SVZ-EGFRwt/amp and SVZ-EGFRvIII tumors were surgically excised from nude mice and the tissue was dissociated of enzymatic digestion at room temperature with Accumax by 30 min. Samples then were filtered through 70 μm strainers and collected in staining medium (PBS containing 2% BSA). Live tumors cells were discriminated from dead cells using propidium iodide and GFP+ cells were isolated with BD FACS Cell Sorter. Cells were collected into 2 ml staining medium and were recovered by centrifugation for further analysis.

### Chicken chorioallantoic membrane (CAM) assay

For in vivo evaluation of the angiogenic inductive potential of SVZ derived CM, a CAM assay was performed as previously reported (11). Photographs were taken with a digital camera HD IC80 (Leica, Heidelberg, Germany) and the number of vessels within a 6-mm radius of the scaffold were counted to determine the angiogenic score, using ImageJ software (NIH, USA). In each photograph the diameter of 220 vessels was measured using ImageJ software (NIH, USA).

### Inmunofluorescent (IF) and Inmunohistochemical (IHC) staining and quantification

IF and IHC staining was performed as previously described (6) using primary and secondary antibodies described in Table S2. For quantification, slides were scanned at 63X or 40X magnification. The number of BrdU-positive cells per field was counted with Fiji-ImageJ software and normalized with the total number of cells. To quantify the IgG extravasation Fiji-ImageJ software was used. The signal from the endomucin channel was subtracted from the IgG channel. To quantify delocalized pericytes, the signal from the endomucin channel was subtracted from the αSMA channel. For the quantification of the vasculature, we counted the number of dilated vessels per field and CD34 staining with Fiji-ImageJ software to show the vascular density. For the quantification of the vasculature, we counted the number of dilated vessels per field using CD34 staining. Density measurements of blood vessel density, necrotic area and pericytes coverage, were performed with ImageJ software (http://rsb.info.nih.gov/ij). Furthermore, in case of the necrotic area we used a score to grade the intensity of the quantified necrosis. To calculate vasculature per random field areas was measured in the intratumoral regions of tumor sections.

### In silico analysis

The Cancer Genome Atlas (TCGA) GBM, LGG and GBM+LGG dataset was accessed via cBioPortal (https://www.cbioportal.org/), UCSC xena-browser (https://xenabrowser.net) and Gliovis (http://gliovis.bioinfo.cnio.es) for extraction of the data: overall survival, gene’s expression level and the distribution of the different genetic alterations. Kaplan-Meier survival curves were done upon stratification based into low and high groups using expression values from each gene. Significance of differences in survival between groups was calculated using the log-rank test. For the functionality studies we have used “David Gene ontolog “analysis. First, we selected a cluster of 365 genes co-expressed with *SOX9*, using the highest value of the spearman’s correlations. Then, “David gene ontology “analysis associates the expression of this genes with the biological processes involved. The hypoxic-related genes signature included hypoxia and HIF1α pathways genes. Ivy Gap date set analysis (http://glioblastoma.alleninstitute.org/) was used to analyze gene signature enrichment between the different anatomic structures identified in the tumor.

### Statistics

GraphPad Prism 5 software was used for data presentation and statistical analysis. For bar graphs, the level of significance was determined by a two-tailed un-paired Student’s t-test. The difference between experimental groups was assessed by Paired t-Test and one-way ANOVA. For Kaplan-Meier survival curves, the level of significance was determined by the two-tailed log-rank test. P values < 0.05 were considered significant (*p < 0.05; **p < 0.01; *** p< 0.001; **** p< 0.0001; n.s., non-significant). All quantitative data presented are the mean ± SEM. Precise experimental details (number of animals or cells and experimental replicates) are provided in the figure legends.

Additional details are provided in the Supplementary Methods.

## Results

### Correlation of IDH1/2 and EGFR genetic alterations with the expression of angiogenic molecules

We first validated that the transcription of three recognized angiogenic markers (*VEGFA, ANGPT2* and *IGFBP2*) shows a strong inverse correlation with the survival of glioma patients (**Figure 1A-C**), even if we consider LGG (Figure S1A-C) and GBM (Figure S1D-F) separately. Moreover, the progressive increase in their expression paralleled the evolution of the glioma disease (**Figure 1D-F**). We then measured the frequency of mutations (**Figure 1G-I**) and copy number amplifications (CNA) (**Figure 1J-L**) in the groups with high or low quantities of *VEGFA* (**Figure 1G-J**), *ANGPT2* (**Figure 1H-K**) and *IGFBP2* (**Figure 1I-L**). As expected, the frequency of *IDH1* mutations was much higher in the gliomas that contain less angiogenic-related mRNAs. Among the rest of the genes analyzed, we found a consistent enrichment in mutations and CNA of *EGFR* and *EGFR*-related molecules, like *ELDR* (EGFR long non-coding downstream RNA) or *EGFR-AS1*, in the most angiogenic tumors.

**Figure 1.**
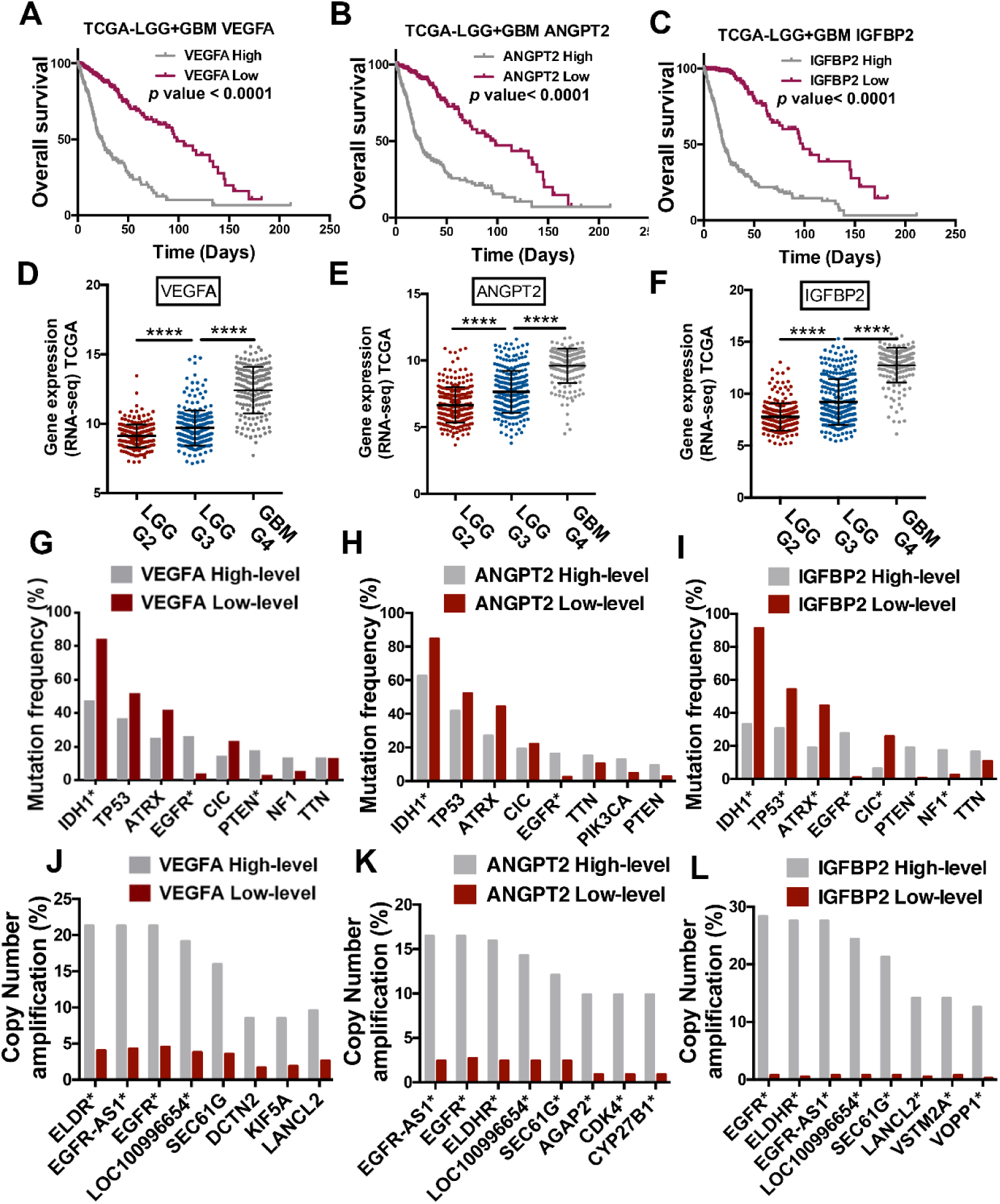
Correlation of EGFR alterations with the expression of angiogenic molecules. **A-C**. Kaplan-Meier overall survival curves of patients (TCGA, LGG+GBM cohort) (n=663), stratified into 2 groups based on high and low *VEGFA* (A), *ANGPT2* (B) or *IGFBP2* (C) expression values. **D-F**. Analysis of *VEGFA* (D), *ANGPT2* (E) and *IGFBP2* (F) expression by RNAseq in gliomas (TCGA, LGG+GBM cohort) (n=663), grouped according to the clinical evolution (grade) of the tumors. **G-L**. Frequency of gene mutations (G-I) or copy number amplification (J-L) in gliomas with high or low level of expression of *VEGFA* (G), *ANGPT2* (H) and *IGFBP2* (I) (n=661). *P ≤ 0.05, ****P ≤ 0.0001.

### Stratification of gliomas by the genetic status of IDH-EGFR distinguishes between three different vascular phenotypes

Tumors with alterations in *IDH1/2* and *EGFR* account for almost 90% of all gliomas in the cancer genome atlas (TCGA) cohort, and they show a mutually exclusive pattern, with only a small percentage of IDHmut gliomas harboring *EGFR* gains (**Figure 2A**). We classified our own cohort of patient’s samples in three groups, IDHmut gliomas, IDHwt gliomas without EGFR mutations (EGFRwt/amp) and IDHwt/EGFRmut gliomas, and we measured the amount of the angiogenic-related mRNAs. As it has been previously described (12), the vasculature of IDHmut tumors was close to normal and there was no overexpression of any of the three angiogenic and vascular markers in these tumors compared to normal brain tissue (**Figure 2B-G**). Among gliomas, we noticed that there was a gradual increase in the amount of the three mRNAs from the IDHmut to the IDHwt/EGFRmut gliomas, being the IDHwt/EGFRwt/amp group in the middle (**Figure 2B-D**). This result was consistent with the in silico data (Figure S2A-C) and with the immunohistochemical (IHC) (**Figure 2E**) and the transcriptional (**Figure 2F-G**, and Figure 2SD-E) analysis of CD34 (ECs) and αSMA (pericytes), which showed a gradual increase in the three glioma subgroups. The parameters that changed the most were the vascular density (**Figure 2H**) and the number of dilated BVs (**Figure 2I**), typical of malignant gliomas (13).

**Figure 2.**
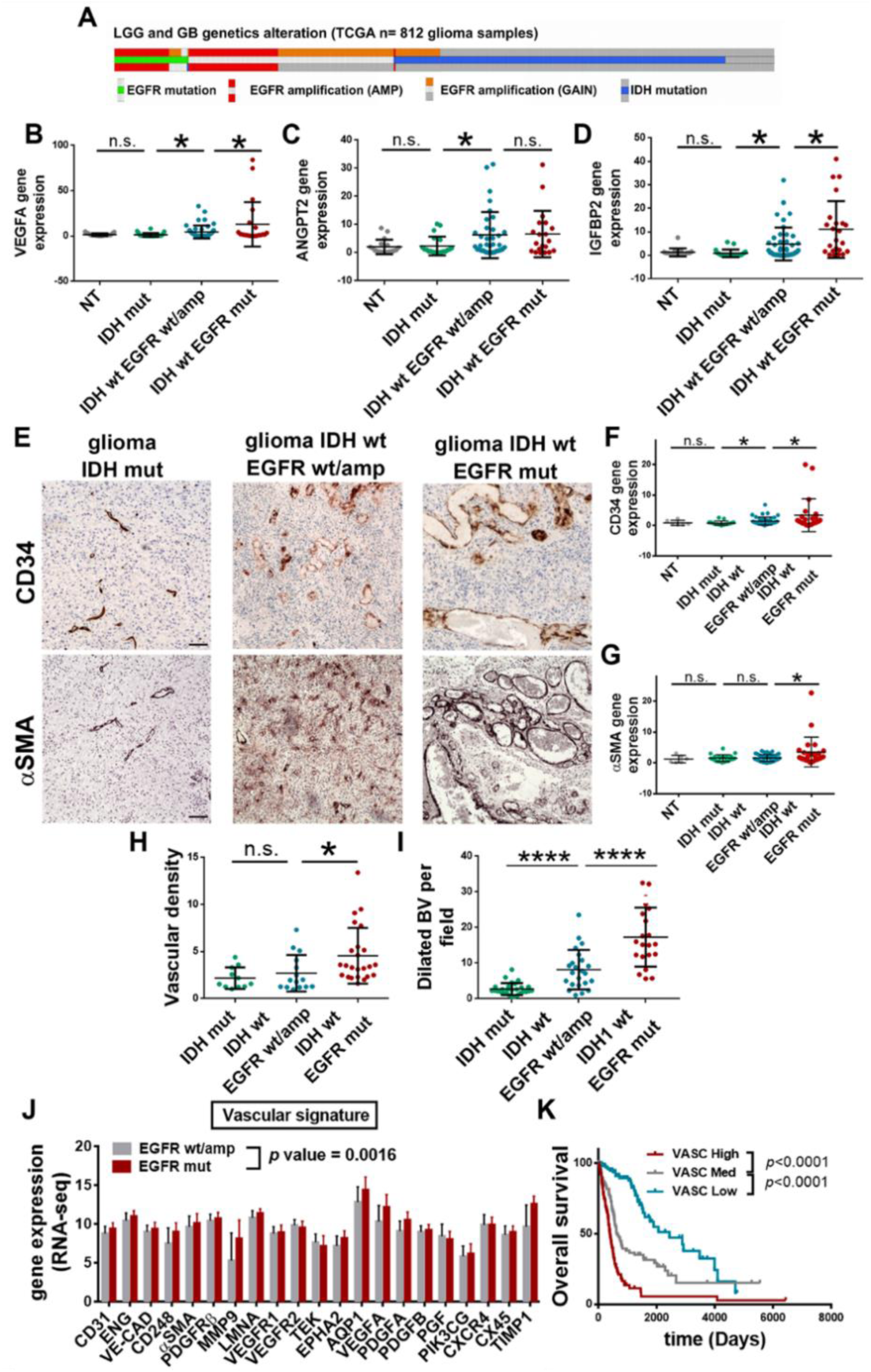
Stratification of gliomas by the genetic status of *IDH-EGFR* distinguishes between different vascular phenotypes. **A**. Distribution of somatic nonsilent mutations in *IDH1/2* and *EGFR* in a TCGA (LGG+GBM) cohort (n=663). **B-D**. qRT-PCR analysis of *VEGFA* (B), *ANGPT2* (C) and *IGFBP2* (D) expression in gliomas (n=93) classified as IDHmut, IDHwt/EGFRwt/amp and IDHwt/EGFRmut. *HPRT* was used for normalization. **E**. Representative pictures of IHC staining of CD34 (Top) and αSMA (Bottom) in three representative tumors. **F-G**. qRT-PCR analysis of *CD34* (F) and *αSMA* (G) expression in gliomas (n=93). *HPRT* was used for normalization. **H**. Quantification of the vascular density in (E) (n=46). **I**. Quantification of the dilated blood vessels (BVs) in (E) (n=46). **J**. RNAseq analysis of angiogenesis-related genes in a TCGA (LGG+GBM) cohort (n=319). **K**. Kaplan-Meier overall survival curves of patients from the TCGA (LGG+GBM) cohort (n=319), stratified into 3 groups based on the expression of the vascular signature. *P ≤ 0.05; ****P ≤ 0.0001. n.s. not significant. Scale bars: 100 μm.

Our results suggest that the *IDH-EGFR*-based stratification of gliomas allows the distinction of three different vascular phenotypes, highlighting important differences between gliomas that overexpress wt or mut EGFR, probably related to the increased phosphorylation of the receptor (Figure S2F), and the increase in the expression of several angiogenesis-related signatures (Figure S2G-J) observed in the second group. We narrowed down the set of differentially expressed vascular genes to define a “vascular signature “and we confirmed its higher expression in EGFRmut vs EGFRamp glioma patients (**Figure 2J**). Moreover, this vascular signature showed a strong correlation with the overall survival of glioma patients (**Figure 2K**).

### EGFR wt/amp and EGFR vIII cells have different vascular capacities

To test the effect of wt or mut EGFR cells in the surrounding vessels, in the absence of any masking effect of concomitant mutations in other genes, we generated two mouse glioma models by transforming subventricular zone (SVZ) progenitors from p16/p19 ko mice with retrovirus carrying the wt or the vIII isoform of the receptor (Figure S3A-B). These cells grew in vitro (Figure S3C) and they were both very sensitive to dacomitinib (Figure S3D), an inhibitor of the receptor’s tyrosine kinase activity (10). The two mouse cell lines formed subcutaneous tumors, although SVZ-EGFRvIII cells grew faster (Figure S3E). Moreover, they were very sensitive to dacomitinib in vivo (Figure 3F-G).

**Figure 3.**
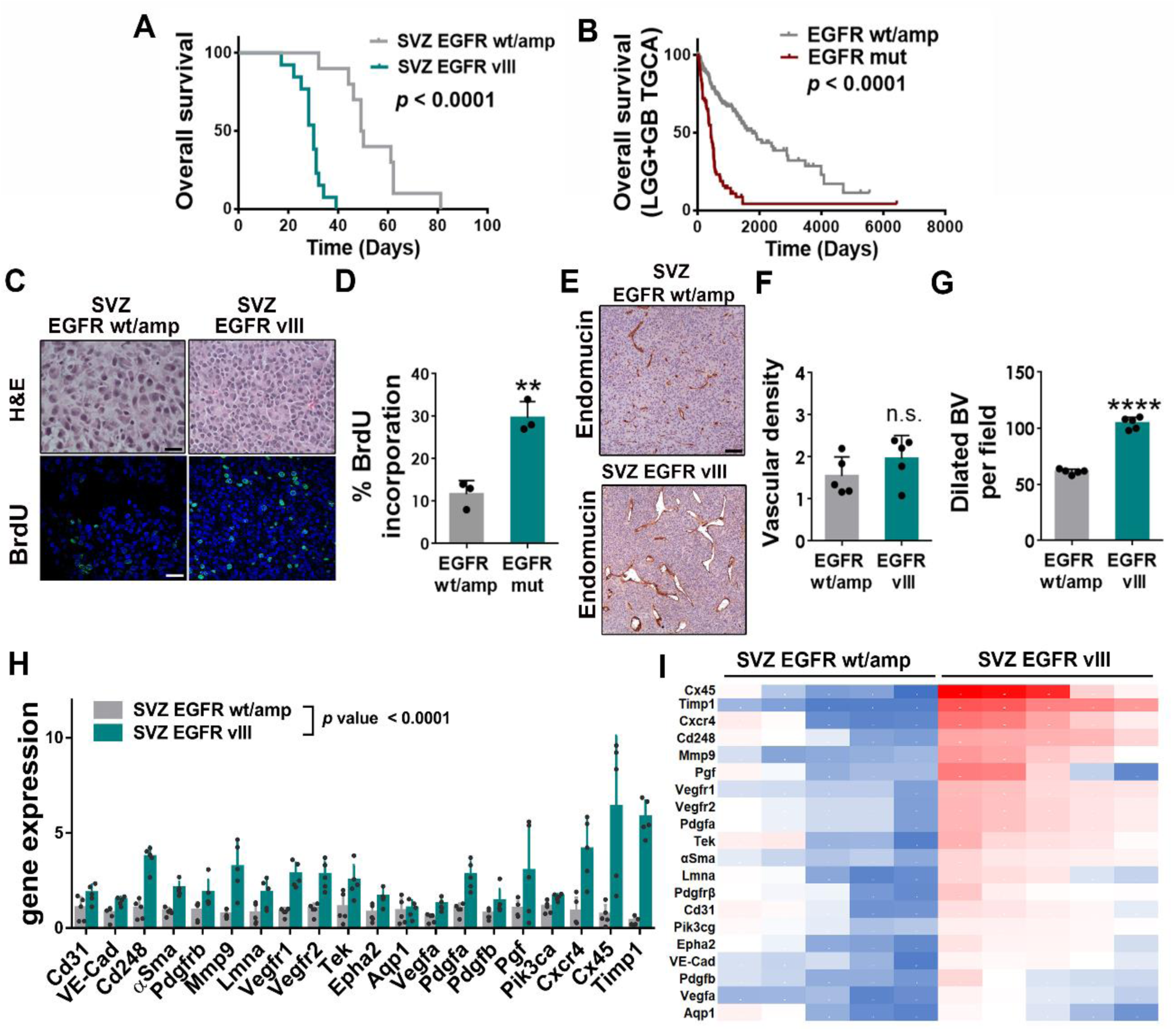
EGFR wt/amp and EGFR vIII cells have different vascular capacities. **A**. Kaplan-Meier overall survival curves of mice that were orthotopically injected with SVZ-EGFRwt/amp or SVZ-EGFRvIII cells (n=10). **B**. Kaplan-Meier overall survival curves of patients from the TCGA cohort (GBM+LGG) separated based on the genetic status of *EGFR* (n=272). **C**. Representative images of hematoxylin and eosin (H&E) (Top) and BrdU uptake (Bottom) in sections from SVZ tumors. **D**. Quantification of the percentage of BrdU+ cells in SVZ tumors (n=3). **E**. Representative images of the endomucin IHC staining of SVZ glioma sections. **F-G**. Quantification of the vascular density (F) and the number of dilated blood vessels (BVs) (G) in (E) (n=5). **H**. qRT-PCR analysis of angiogenesis-related genes in SVZ-EGFRwt/amp and SVZ-EGFRvIII tumors. *Actin* was used for normalization (n=5). **I**. Heat map of gene expression analysis in H. Red: highest expression. Blue: lowest expression. ** P ≤0.01; ****P ≤0.0001; n.s. not significant; Scale bars: 25 μm (C), 100 μm (E).

Orthotopic implantation of the SVZ models generated gliomas with a high penetrance. Tumor burden was higher after the injection of vIII-expressing cells (**Figure 3A**), which correlates with the worse prognosis of EGFRmut gliomas in comparison with those harboring the wt receptor (amplified or not) (**Figure 3B**). The IHC and the immunofluorescent (IF) analysis of the tumors revealed a higher compact and proliferative growth after EGFRvIII expression (**Figure 3C-D**). The vascular density was not significantly different between the two models (**Figure 3E-F**) but we observed a strong increase in the size of the vessels in EGFRvIII gliomas (**Figure 3E-G**).

To obtain an independent confirmation of this observation, we used the chicken embryo chorioallantoic membrane (CAM) assay, which has been widely used to study angiogenesis (14). Bio-cellulose scaffolds were embedded in VEGFA (as a control) or conditioned media (CM) from SVZ-EGFRwt/amp or SVZ-EGFRvIII cells, and they were layered on top of growing CAMs (Figure S3H). Although the CM from both type of cells demonstrated a pro-angiogeneic capacity (Figure S3I), the diameters of the vessels formed were larger in the presence of CM from SVZ-EGFRvIII cells compared to their wild-type counterparts (Figure S3J). Moreover, we observed that the vascular signature was strongly upregulated in SVZ-EGFRvIII compared to SVZ-EGFRwt/amp tumors (**Figure 3H-I**). Notably, some of the top genes in this comparative analysis were linked to pericytic differentiation (*Cd248*) and function (*Timp1*) (**Figure 3I**). Overall, these results confirm the pro-angiogenic function of EGFR signaling in gliomas. Moreover, they suggest that the expression of different isoforms of the receptor have a distinct effect on the vascular microenvironment.

### EGFRwt/amp expression is associated with a hypoxic phenotype

To our surprise, the magnetic resonance imaging (MRI) analysis of the grafted mice revealed a cumulative increase in the contrast enhancement of SVZ-EGFRwt/amp compared to SVZ-EGFRvIII injected brains (**Figure 4A-B**). We also observed a stronger extravasation of Evans Blue (Figure S4A) and IgG (**Figure 4C-D**) in SVZ-EGFRwt/amp compared to mutant tumors. These observations suggest that the integrity of the BBB is severely compromised in the former. The disruption of the BBB has been associated with impaired blood perfusion and the formation of hypoxic regions in tumors (15). Accordingly, HIF1α expression (**Figure 4E**) and the hypoxia-related signature (**Figure 4F**) were higher in SVZ-EGFRwt/amp compared to SVZ-EGFRvIII tumors. This signature was defined using the IvyGAP (Ivy GBM Atlas Project) data-set analysis (Figure S4B), selecting the most relevant genes included in hypoxia and HIF1α pathways signatures that were up-regulated in the perinecrotic and pseudopalisading-cell-necrosis tumor zones (Figure S4C). Notably, this signature was increased in mesenchymal (MES), compared to classic (CL) or proneural (PN) gliomas (Figure S4D) (16). MES gliomas are characterized by a high frequency of EGFR amplifications (but not mutations) and show a higher overall fraction of necrotic area and a stronger expression of hypoxia-regulated genes compared to the other subtypes (16, 17). By contrast, tumors harboring EGFR mutations tend to accumulate in the CL subgroup, characterized by a highly proliferative phenotype (16, 18).

**Figure 4.**
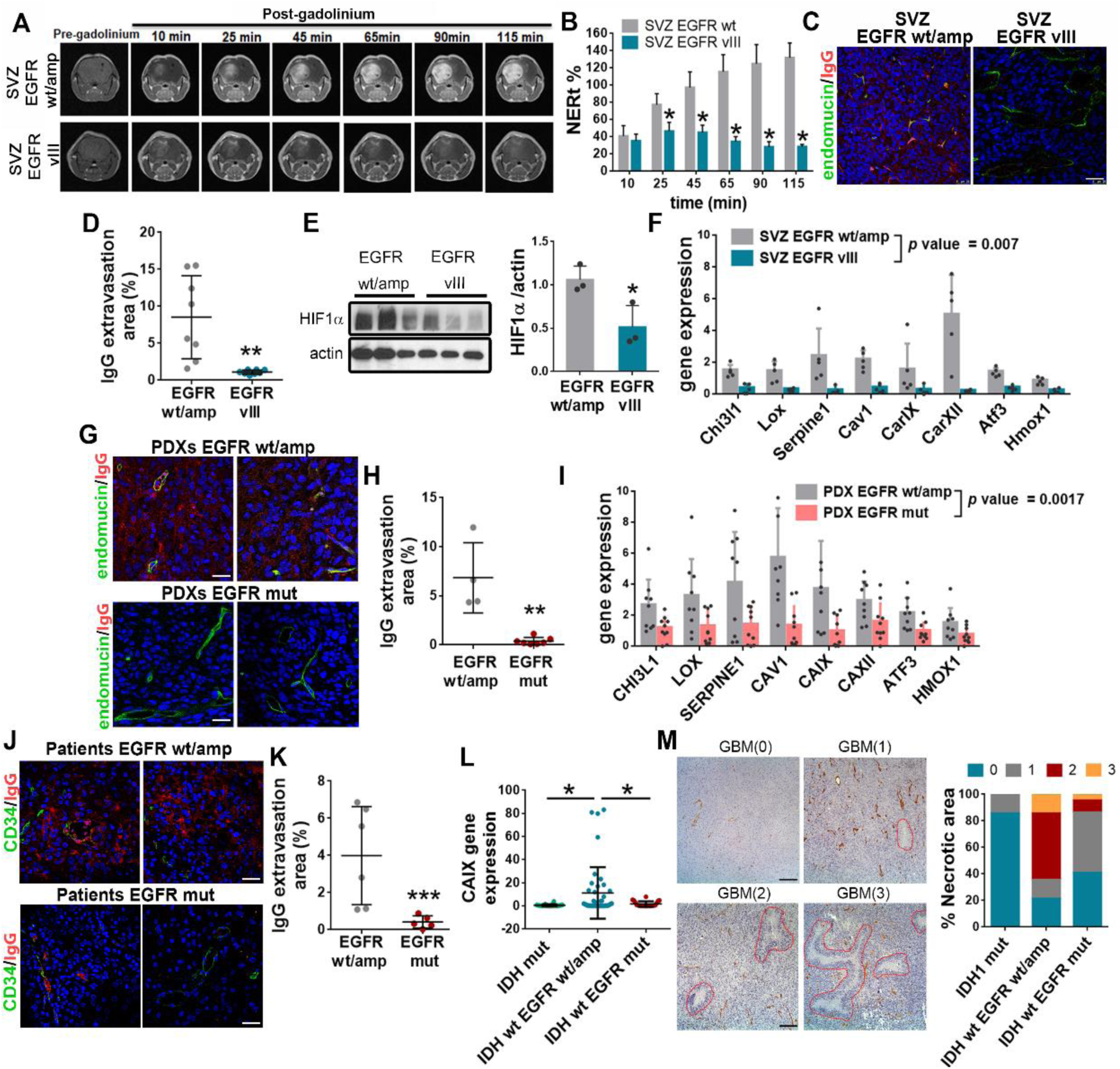
EGFRwt/amp expression is associated with a hypoxic phenotype. **A**. Representative T1 contrast enhanced MRI scans of mouse brains containing SVZ tumors at different time points after gadolinium (Gd) injection. **B**. Quantification of the Gd extravasation in (A) (n=3). NERt: normalized enhancement ratio. **C**. Representative images of endomucin and IgG IF staining of sections from SVZ tumors. **D**. Quantification of IgG extravasation on (C) (n=16). **E**. WB analysis and quantification of HIF1α in SVZ tumors. Actin was used for normalization. **F**. qRT-PCR analysis of hypoxic-related genes signature in SVZ tumors (n=5). *Actin* was used for normalization. **G-H**. Representative images (G) and quantification (H) of endomucin and IgG IF staining of sections from PDXs (n=11). **I**. qRT-PCR analysis of hypoxic-related genes signature in EGFRwt/amp (n=9) and EGFRmut (n=10) PDXs. *HPRT* was used for normalization (PDXs). **J**. Representative images of endomucin and IgG IF staining of sections from human glioma samples. **K**. Quantification of IgG extravasation on (J) (n=10). **L**. qRT-PCR analysis of *CAIX* expression in human glioma samples. *HPRT* was used for normalization (n=89). **M**. Representative pictures of the hematoxylin and eosin (H&E) staining of different GBMs. Necrotic areas are highlighted with a red line and the necrotic area score is represented between brackets. The percentage of tumors (n=48) with different necrotic area scores is shown on the right. *P ≤ 0.05; **P ≤ 0.01; ***P ≤ 0.001; Scale bars: 50 μm (C), 25 μm (G, J), 100 μm (M).

We then performed an IF analysis in different orthotopic patient-derived-xenografts (PDXs) that express EGFRwt/amp or EGFRmut (deletions and/or point mutations). We observed the increase in the permeability of IgG in the prior compared to the latter (**Figure 4G-H**). This effect was paralleled by an upregulation of the hypoxia-related signature in EGFRwt/amp PDXs (**Figure 4I**). Notably, the extravasation of IgG was also increased in EGFRwt/amp compared to EGFRmut patients’ tumors (**Figure 4J-K**). Besides, we found a strong increase in the expression of *Carbonic Anhydrase IX (CAIX)*, one of the main HIF1α targets, in IDHwt/EGFRwt/amp gliomas compared to tumors with *IDH* or *EGFR* mutations (**Figure 4L**). IDHmut tumors contain a more “normalized “vasculature without vessel leakage (6) and a reduced extent of necrosis (19). The histological analysis of the tumor’s sections confirmed these observations (**Figure 4M**). Moreover, it showed an accumulation of necrotic areas in EGFRwt/amp compared to EGFRmut gliomas (**Figure 4M**). These results suggest that the angiogenic signals induced by amplification or overexpression of wt EGFR result in a dense vascular network, but with a severely compromised BBB, a different phenotype to the one observed in mutant EGFR tumors. This vascular fragility is associated with a less efficient fueling of tumor proliferation and with the appearance of areas of necrosis and hypoxia.

### Glioma derived-pericytes stabilize the vasculature in EGFRmut tumors

SVZ-EGFRvIII tumors showed a highly compact growth, whereas in SVZ-EGFRwt/amp tumors cells appeared detached from each other and from the tumor vessels (**Figure 5A**). It has been shown that pericytic functions can be performed by the highly plastic glioma stem cells, which acquire mesenchymal and mural cell features (3) in a process regulated by EGFR signaling (6). In agreement with these notions, some tumor cells expressed αSMA in both SVZ glioma models (**Figure 5B**). Furthermore, pericyte-related markers were expressed in GFP+ cells, sorted out after tumor dissociation, independently of the genetic status of *EGFR* (**Figure 5C**). However, we noticed an increased transcription of *αSma* and *Cd248* in SVZ-EGFRvIII tumor cells, compared to their wild-type counterparts (**Figure 5D**). Notably, among the vascular signature, several pericyte-related genes were upregulated in cultured SVZ-EGFRvIII cells (Figure S5A), suggesting that they express these markers even in the absence of the microenvironment-derived signals, and that this process is exacerbated in the presence of the mutant isoforms of the receptor. In agreement with these results, there was a significant increase in the expression of human *αSMA* and *CD248* (**Figure 5E**) and in the ratio of tumor pericytes to mouse ECs (Figure S5B) in EGFRmut compared to EGFRwt/amp PDXs, whereas the transcription of host pericyte genes in all the PDX tested was similar to that of normal mouse brain (**Figure 5E**, and Figure S5B). Notably, human endothelial transcripts (*CD31, END*) were not overexpressed in the PDXs, independently of the genetic status of *EGFR* (Figure S5C). These results suggest that in the presence of EGFR mutations glioma cells have a higher capacity to differentiate into pericytes but not to ECs. Indeed, we found that up to 80% of the αSMA+ cells in a patient’s tumor express the vIII mutation (Figure S5D). Moreover, we found an increased pericyte coverage in mouse (**Figure 5F**) and human (**Figure 5G**) glioma transplants. By contrast, we noticed that the amount of delocalized pericytes (those that were not in close contact with ECs) was higher in wild-type compared to mutant EGFR tumors (Figure S5E-F).

**Figure 5.**
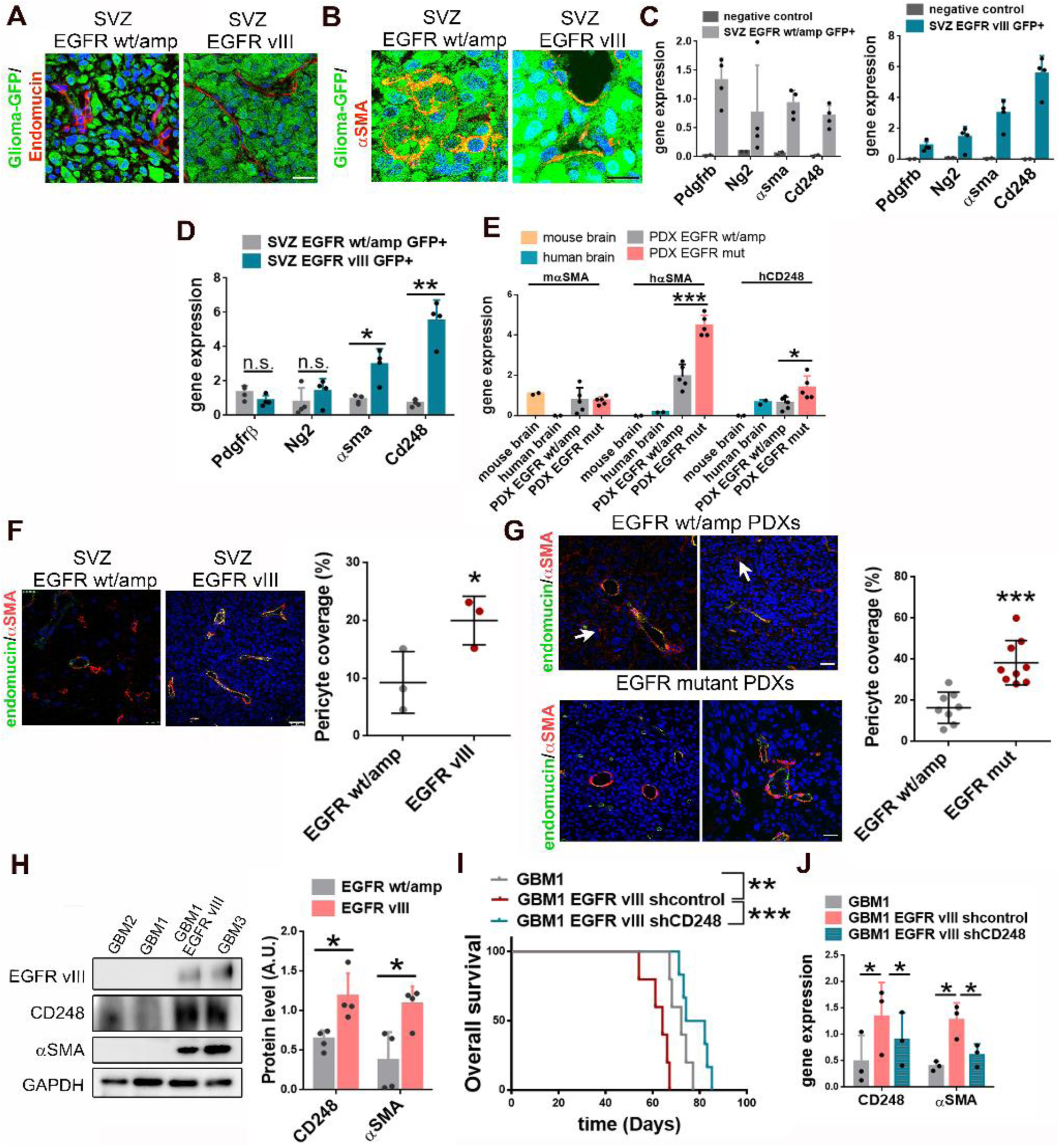
Glioma derived-pericytes stabilize the vasculature in EGFRmut tumors. **A-B**. Representative pictures of GFP+ glioma cells and endomucin (A) or αSMA (B) IF staining of sections from SVZ tumors. **C**. qRT-PCR analysis of pericytic-related genes in GFP+ sorted cells from SVZ tumors (n=4). Human cDNA was used as negative control. *Actin* was used for normalization. **D**. Comparative analysis of expression of pericytic markers in tumors from (C). **E**. qRT-PCR analysis of pericytic–related genes in EGFRwt/amp and EGFRmut PDXs (n=5). Human and mouse tissue were used as control. *HPRT* or *Actin* were used for normalization. **F-G**. Representative images of endomucin and αSMA IF staining of sections from SVZ gliomas (F) and PDXs (G). Arrows point towards examples of αSMA-positive cell that do not localize close to endomucin+ cells. Quantification of the pericyte coverage is shown on the right of the images (F, n=10). (G, n=8). **H**. WB and quantification of EGFRvIII, αSMA and CD248 in GBM1 and GBM2 (EGFRwt/amp), GBM1-EGFRvIII, and GBM3 (EGFRamp/mut) cells. GAPDH was used as loading control. **I**. Kaplan-Meier overall survival curves of mice that were orthotopically injected with GBM1, GBM1-EGFRvIII-shcontrol and GBM1-EGFRvIII-shCD248 (n=6). **J**. qRT-PCR analysis of *CD248* and *αSMA* in the tumors in (I). *HPRT* was used for normalization (n=3). **\***P ≤ 0.05; **P ≤ 0.01; ***P ≤ 0.001; n.s. not significant. Scale bars: 10μm (A-B), 25μm (F-G).

To further study the vascular role of mutant EGFR, we introduced EGFRvIII in GBM1 cells (EGFRamp). We observed a strong increase in the amount of αSMA and CD248 protein in vitro, similar to the expression observed in GBM3 cells (EGFRamp/EGFRvIII) (**Figure 5H**). Overexpression of the mutant receptor increased the aggressiveness of the GBM1 tumors (**Figure 5I**) and induced a higher transcription of human-specific pericyte genes (**Figure 5J**). Notably, downregulation of *CD248*, a master regulator of pericyte differentiation in malignant solid tumors (20, 21), decreased the aggressiveness of GBM1-EGFRvIII tumors (**Figure 5I**), with a concomitant decrease in the expression of other pericyte makers (**Figure 5J**). Taken together, these results confirm that EGFR mutations promote the growth of gliomas, at least in part through the increase in the plasticity of the tumor cells, which can work as pericytes and reinforce the stability of the vessels.

### EGFR mutations modulate the vascular properties of glioma cells in a BMX- and SOX9-dependent way

EGFR signaling activates transcription factors (TFs) that drive tumor growth. We performed an in silico analysis in order to find TFs overexpressed in EGFRmut gliomas but with a similar expression in wild-type or amplified *EGFR* gliomas. The best hit was the Sex-determining region Y (SRY)-box 9 (*SOX9*) gene (Figure S6A), which have been previously linked to EGFR (22, 23). We found a strong upregulation of the SOX9 protein (**Figure 6A**) and mRNA (**Figure 6B**) in those PDXs harboring *EGFR* mutations. Similarly, SOX9 was accumulated in SVZ-EGFRvIII allografts, but not in SVZ-EGFRwt/amp or in GL261 (commonly used mouse glioma cells) tumors (Figure S6B). Furthermore, in the presence of a proteosomal inhibitor the amount of SOX9 was increased in EGFRwt/amp cells but did not change in EGFRmut cells (**Figure 6C**), suggesting the stabilization of the protein in the latter.

**Figure 6.**
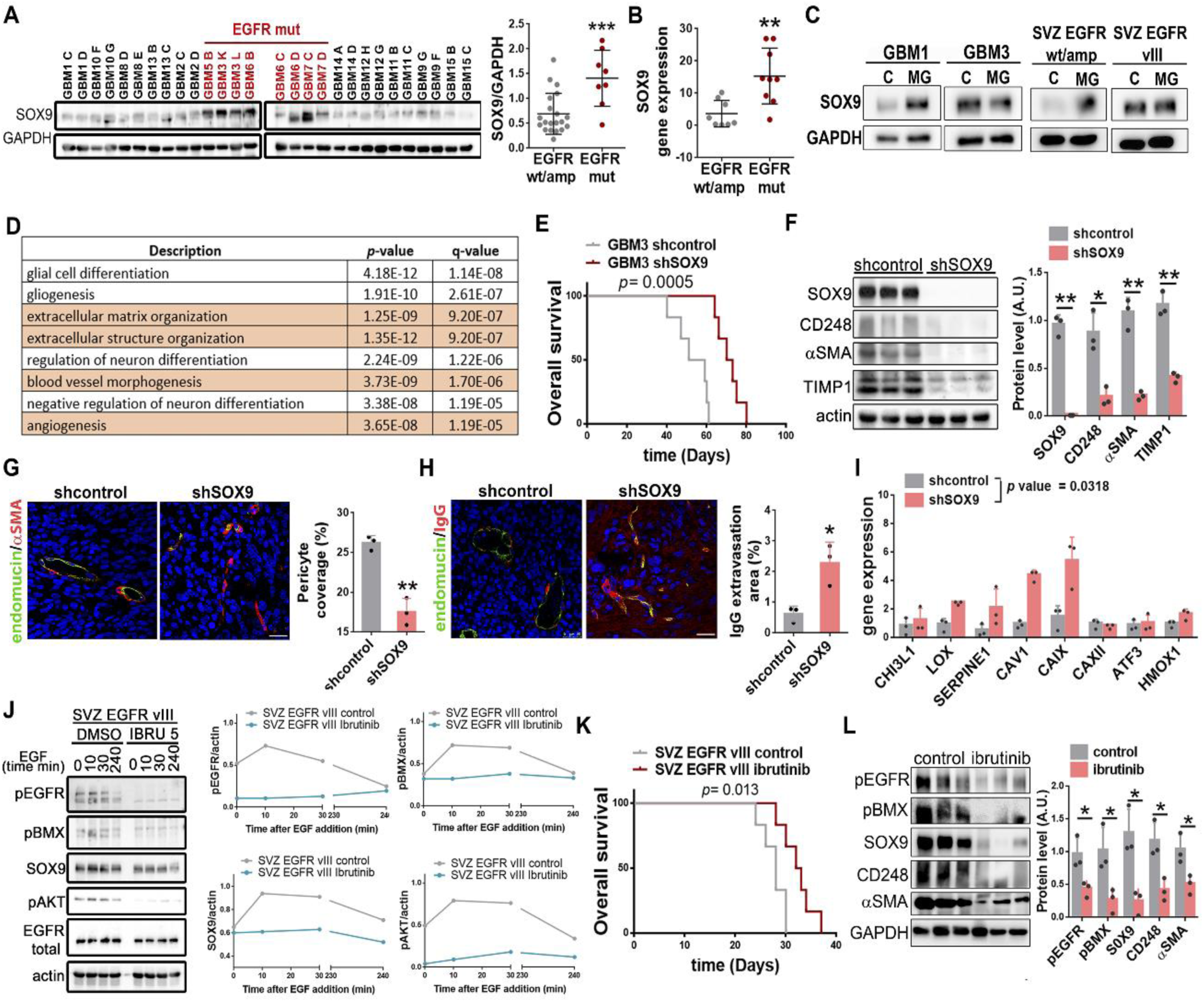
EGFR mutations modulate the vascular properties of glioma cells in a BMX- and SOX9-dependent way. **A**. WB analysis and quantification of SOX9 expression in PDXs. GAPDH was used for normalization. **B**. qRT-PCR analysis of *SOX9* expression in PDXs (n=9). *HPRT* expression was used for normalization. **C**. WB analysis of SOX9 in EGFRwt/amp cell lines (GBM1 and SVZ) and EGFRvIII cell lines (GBM3 and SVZ) in the absence or in the presence of MG132 (MG) (10µm). **D**. David Gene Ontology Enrichment Analysis performed on the genes that were co-expressed with *SOX9* (TCGA (LGG+GB) cohort). **E**. Kaplan-Meier overall survival curves of mice that were orthotopically injected with GBM3-shcontrol or GBM3-shSOX9 cells (n=6). **F**. WB analysis and quantification of SOX9, CD248, αSMA and TIMP1 in the tumors in (E) (n=3). Actin was used as loading control. **G**. Representative images of endomucin and αSMA IF staining of tumors in (E). Quantification of the pericyte coverage is shown on the right (n=3). **H**. Representative images of endomucin and IgG IF staining of sections from tumors in (E). Quantification of IgG extravasation is shown on the right (n=3). **I**. qRT-PCR analysis of hypoxic-related genes in the tumors in (E) (n=3). *HPRT* was used for normalization. **J**. WB analysis and quantification of pEGFR, pBMX, SOX9, pAKT and total EGFR in growth factor-starved SVZ-EGFRvIII cells incubated with EGF (100ng/ml) for the times indicated, in the presence of DMSO or Ibrutinib (5µm). Actin was used for normalization. **K**. Kaplan-Meier overall survival curves of mice that were orthotopically injected with SVZ-EGFRvIII cells and treated with intraperitoneal injections of ibrutinib (12mg/kg/day) (n=6). **L**. WB analysis and quantification of pEGFR, pBMX, SOX9, CD248 and αSMA in the tumors in (K). GAPDH was used for normalization. **\***P ≤ 0.05; **P ≤ 0.01; ***P ≤ 0.001; ****P ≤ 0.0001. Scale bars: 25 μm (G-H).

SOX9 participates in a variety of functions during development, although it has also been implicated in the regulation of cancer stem cells (24). Moreover, we found a positive association of *SOX9* with angiogenic processes in gliomas (**Figure 6D**). Downregulation of this protein in GBM3 (EGFRamp/EGFRvIII) (**Figure 6E-6F**) or in GBM1-EGFRvIII (Figure S6C-D) cells impaired their orthotopic growth. Notably, the expression of *CD248* and other pericyte-related genes was decreased in GBM3-shSOX9 compared to control tumors (**Figure 6F**, and Figure S6E), suggesting that SOX9 mediates the induction of pericyte properties in EGFRmut tumors. In agreement with that, we found a positive correlation between the transcription of *SOX9* and different pericytic-related genes in gliomas (Figure S6F-I). Notably, we observed a decrease in the pericyte coverage (**Figure 6G**), concomitant with the upregulation of the IgG extravasation (**Figure 6H**) and the hypoxia-related signature expression (**Figure 6I**) in *SOX9* interfered tumors. These observations reinforce the idea that blockade of the transdifferentiation capacity of EGFRmut GBM cells favors the fragility of the tumor vessels and the subsequent induction of hypoxia, similar to what occurs in tumors over-expressing wild-type EGFR.

A recent study has discovered that the bone marrow and X-linked (BMX) nonreceptor tyrosine kinase is highly expressed in glioma derived-pericytes but not in normal mural cells in the brain (25). However, the upstream signals have not been described yet. In order to test if EGFR signaling could be connected to BMX activation in glioma cells, we performed an in vitro analysis in response to EGF. Stimulation with the ligand induced EGFR activation and signaling in SVZ-EGFRvIII (**Figure 6J**) and SVZ-EGFRwt/amp (Figure S6J) cells. However, it only stimulated BMX phosphorylation in the presence of the mutation (**Figure 6J-J**). Moreover, we observed an accumulation of SOX9 protein at short times after EGF stimulation in EGFRvIII (**Figure 6J**) but not in EGFRwt/amp cells (Figure S6J). EGF stimulation in GBM7 cells (EGFRamp/EGFRmut), induced a strong BMX activation and the accumulation of SOX9 (Figure S6K), although the time-pattern was different from the one observed in vIII cells. Ibrutinib, a dual BMX/BTK (Bruton’s tyrosine kinase) inhibitor that impairs tumor-to-pericyte transdifferentiation (25), blocked BMX stimulation and SOX9 accumulation in response to EGF in SVZ-EGFRvIII (**Figure 6J**) and GBM7 (Figure S6K) cells. EGFR and AKT phosphorylation were also impaired in the presence of ibrutinib (**Figure 6J**, and Figure S6J). Furthermore, ibrutinib impaired tumor growth of SVZ-EGFRvIII cells (**Figure 6K**) and reduced the phosphorylation of EGFRvIII and BMX in the treated tumors (**Figure 6L**). These changes were paralleled by a reduction in the expression of *SOX9* and pericyte markers (**Figure 6L**). These results suggest that EGFRmut-BMX signaling induces the accumulation of SOX9 protein and the subsequent formation of tumor-derived-pericytes, increasing the aggressiveness of EGFRmut gliomas but rendering these tumors sensitive to ibrutinib.

## Discussion

The results presented here stablish a positive correlation of *EGFR* genetic alterations with the expression of angiogenic molecules and with the appearance of vascular abnormalities, which were absent in the less aggressive IDHmut gliomas. These vascular effects were increased in mutant compared to wt/amp EGFR tumors, especially the presence of dilated BVs. EGFR activity have been previously associated with the angiogenic properties of aggressive gliomas (8, 26, 27). However, our data suggest for the first time that the overexpression of wt or mutant EGFR in glioma cells induce the formation of two different vascular phenotypes, which are distinct from the normalized vasculature of mutant IDH1/2 tumors. The first one is characterized by the instability of the BBB and the induction of hypoxic signals and necrotic areas in the tumors, whereas the second phenotype, induced by EGFR mutations, contains more robust and enlarged tumor vessels that nurture a very compact and hyperproliferative tumor tissue. In agreement with this, the MRI analysis of the SVZ mouse models showed an accumulation of Gd over time in the absence of EGFR mutations. These results could seem contradictory with the literature as it has been suggested that EGFRvIII gliomas have a higher perfusion values of the transfer coefficient (Ktrans) in T1-weighted dynamic contrast-enhanced (DCE)-MRI analyses (28). Similar results were obtained in rat (29) and mouse (26) models. This parameter is related to the degree of vessel permeability, but it is also influenced by the blood flow and the vessel area, which are enlarged in SVZ-EGFRvIII compared to SVZ-EGFRwt/amp tumors. In conventional MRI sequences, measured at short times after contrast administration, Gd extravasation could be higher in EGFR mutant tumors due to the increased blood flow. However, at longer times after Gd injection, EGFRwt/amp tumors would show more contrast enhancement due to the presence of leakier tumor vessels. Although long MRI studies are non-viable in clinical practice, emerging evidence indicates that multiparametric analysis of DCE-MRI data could offer greater insight into the status of the tumor vasculature, especially when measuring the response to anti-angiogenic therapies (30). Using such an approach, no major differences were described in the Gd enhancement of EGFRvIII gliomas, which were otherwise characterized by increased cell density and blood flow, compared to other tumors (31). A similar study has revealed the existence of two different subtypes of IDHwt GBMs: a glycolytic phenotype with predominant neovasculature and a necrotic/hypoxic dominated phenotype. However, no correlation was stablished with the genetic status of *EGFR* (32). These kind of studies or other approaches to measure tumor vessel caliber and/or structure in gliomas (13) would certainly help to characterize the two different vascular ecosystems in IDHwt GBMs and how they evolve in response to therapy.

Our data shows an induction of HIF1α protein and function in EGFRwt/amp compared to EGFRmut glioma models. A direct transcriptional activation of *HIF1α* by EGFR has been proposed in lung cancer, especially in the presence of mutations (33), but little is known about this interaction in gliomas. Although we cannot discard a direct regulation of *HIF1α* expression by EGFR activation, we believe that this TF is induced and/or stabilized in the tumor context in response to the environmental conditions (such as acidosis or nutrients/oxygen deprivation) that occur after the disruption of the BBB in EGFRwt/amp gliomas. Notably, there seems to be a reciprocal relation as tumor hypoxia up-regulates *EGFR* expression in lung cancers (33), as well as in gliomas (34), promoting its activation in the absence of mutations. This could serve as a pro-survival signal for hypoxic cancer cells. Moreover, hypoxia has been proposed to induce resistance to EGFR inhibitors in different cancers (35), suggesting that combinatorial approaches targeting both pathways could be a promising strategy for EGFRwt/amp gliomas.

Pericytes play an essential role maintaining the structure and function of the vessel wall and the BBB. These mural cells were thought to originate from mesenchymal progenitors that are recruited from the bone-marrow in response to hypoxia (36, 37). However, it has been recently shown that the majority of vascular pericytes in GBMs derive from tumor cells (3). A similar epithelial-to-pericyte transition has been proposed in other cancers, promoting vascular integrity and tumor growth (38). Nevertheless, to date it was unknown if different genetic alterations could modulate this transdiferentation process. Results from this study and our recently published data (6) suggest that it is governed by EGFR signaling. Moreover, we have shown here that tumor cells expressing mutant isoforms of EGFR have a higher capacity to differentiate into mural cells and overexpress molecules such as CXCR4, CX45 and TIMP1, which could be responsible for the improved pericyte recruitment toward endothelial cells and the increase pericyte coverage observed in EGFRmut gliomas. However, downregulation of *CD248* expression in the tumor cells impaired pericyte differentiation and was sufficient to block the pro-tumorigenic potential of EGFR mutations. This effect was paralleled by a destabilization of the tumor vessels and the increase on hypoxic signals, highlighting the relevance of the tumor-derived pericytes and the angiogenic molecules secreted by them in the regulation of the vascular stability.

The plasticity of the tumor cells in EGFRmut gliomas confers a higher aggressiveness to the tumors. However, it also increases their sensitivity to molecules that target pericytic function, such as ibrutinib. This is especially interesting as it specifically affect tumor-derived pericytes (25) and cancer stem cells (39), preserving normal brain cells, which could limit the toxicity of the drug. Besides, this compound inhibits mutant EGFR activity in gliomas (40), as well as in other cancers (41). Ibrutinib is being tested on unselected GBM patients in combination with radio- and chemotherapy (NCT03535350). We propose that its activity could be higher in tumors harboring EGFR mutations so future retrospective studies should be carried on to validate this idea.

Downstream of EGFRmut/BMX signaling, we have found an enrichment of *SOX9* expression and protein stability. The oncogenic function of SOX9 has been proposed in different cancers (24), including gliomas (42, 43), where it induces proliferation and cell survival, partly through BMI1 upregulation (44). Our data indicate that SOX9 also regulates the vascular properties of gliomas by inducing the cellular plasticity of tumor cells, which could be linked to its well-known effects in cancer stem cells (24). Notably, SOX9 has been linked to the upregulation of PDGFRα expression during development (45), and regulates genes involved in the extracellular matrix such as collagen, aggrecan and Timp1, which are involved in the maintenance of the blood vessel structure (46). In addition, BMI1 has been linked to the regulation of the epithelial-to-mesenchymal transition (47), allowing us to propose that SOX9, either acting directly or through the modulation of BMI1, may act as a master regulator of the vascularization processes in gliomas, especially for those tumors carrying *EGFR* mutations.

In summary, we propose the existence of two IDHwt GBM phenotypes orchestrated by the genetic status of *EGFR* and the downstream modulation of BMX-SOX9 activity, which induces the transdifferentiation of tumor cells into pericytes. This model could have diagnostic as well as great predictive value, as the different subtypes could have a distinct sensitivity to anti-angiogenic or vessel-normalization strategies. Moreover, the function of the tumor-derived-pericytes could limit the entrance of cytotoxic therapies through the BBB (25), which might explain the increased resistance of EGFRmut gliomas. Moreover, the immune infiltrate and its pro- or anti-tumoral properties could also be affected by the vascular abnormalities observed in IDHwt gliomas (48). Overall, our results place the angiogenic properties of gliomas at the top of the microenvironment hierarchy and suggest that future combinatorial therapeutic approaches should combine agents targeting the glioma vasculature with conventional therapies, molecularly-directed drugs or immunotherapies. Moreover, these strategies should be tailor-designed for each specific glioma subtype.

## Availability of data and material

All data associated with this study are present in the paper or in the Supplementary Materials.

## Ethics approval and consent to participate

Glioma tissues were obtained after patient’s written consent and with the approval of the Ethical Committee at Hospital 12 de Octubre (Madrid, Spain) (CEI 14/023 and CEI 18/024).

Animal experiments were reviewed and approved by the Research Ethics and Animal Welfare Committee at our institution (Instituto de Salud Carlos III, Madrid) (PROEX 244/14 and 02/16), in agreement with the European Union and national directives.

## Author’s contributions

Designing research studies: RG, BSC, and PSG; Acquiring data: RG, BSC, MGA, BH, EHS and BC; Analizing data: BSC, RG, VP and PSG; Resources: VP, AM, APN, JMS and AHL; Writing-Original Draft: RG, BSC and PSG; Writing-Review & Editing: MGA, BH, EHS, VP and AM; Funding Acquisition: RG, JMS and PSG; Supervision: RG and PSG.

## Acknowledgments

We would like to acknowledge Manuel Serrano for kindly donating the p16/p19 ko mice and Jacqueline Gutiérrez and Rafael Hortigüela for their technical support. Figure 7 was created with BioRender.

