## Supplementary material for "The genetic status of IDH1/2 and EGFR dictates the vascular landscape and the progression of gliomas"

**Supplementary materials**

Supplementary methods

Table S1. List of GBM cell lines.

Table S2. Antibodies.

Table S3. Primers used for the qRT-PCR analysis.

Supplementary Figures

**Supplementary methods**

*Lentiviral/retroviral production*

Lentiviral and retroviral particles were produced in 293T cells with packaging plasmid pCMVdR8.74 (Addgene #Plasmid 22036) and VSV-G envelope protein plasmid pMD2G (Addgene #Plasmid 12259) using Lipofectamine and Plus reagent (Invitrogen).

*In vitro assays*

For the time-course experiments, cells were starved for two hours in the presence of 5 µM ibrutinib or DMSO before adding 100 ng/µl of EGF. Cells were collected at 0, 10, 30 and 240 min before being chilled on ice and pelleted for WB. For MG132 (Millipore) treatment, 10 µM MG132 was added to the cells and collected 3 h after. Pellets were obtained for WB analysis.

To test the viability of SVZ cells, 10000 cells grown in a 96-well microplate were incubated in the presence of dacomitinib (Pfizer, PF-299804) or vehicle (DMSO) for three days and cell viability was assessed by a colorimetric assay using a WST-1 reagent (Roche) according to manufacturer´s instructions.

*Intracraneal tumor formation*

Intracraneal orthotopic xeno- and allo-grafts were performed using a Hamilton syringe to inject 100.000-300.000 cells (resuspended in 2 µl of stem cell medium) into athymic Nude-Foxn1nu brains (Harlan Iberica). The injections were made into the striatum (coordinates: A–P, −0.5 mm; M–L, +2 mm, D–V, −3 mm; related to Bregma) using a Stoelting Stereotaxic device. When applicable, tumor growth was monitored in an IVIS equipment (Perkin Elmer) after intraperitoneal injection of D-luciferin (75 mg/Kg) (PerkinElmer). The animals were sacrificed at the onset of symptoms.

*Heterotopic allografts*

SVZ-EGFRwt/amp and SVZ-EGFRvIII cells ($1\times 10^6$) were resuspended in culture media and Matrigel (BD) (1:10) and then subcutaneously injected into nude mice. When tumors reached a visible size the tumor volume was measured with a caliper every 5 days. Tumor volume = 1/2(length × width2).

*Mouse magnetic resonance imaging (MRI) and quantification*

Global shimming was first performed and three scout images in axial, sagittal, and coronal direction were acquired using a T1 weighted spin echo sequence with a repetition time of 2.1 s and an effective echo time of 62 ms. The field of view (FOV) was of 3.0x3.0 $cm^2$, the thickness of the slices was 2.0 mm and the matrix size was 256x128, with a total acquisition time of 33 s. Then respiratory-gated T1-weighted spin echo images in coronal orientation were acquired for the tumor visualization with the next parameters: TR/TE = 505/10 ms; FOV=2.56x2.56 $cm^2$; slice thickness: 1.0 mm; number of slices: 9; number of signal averages = 4; matrix size: 256x192. Data were zero filled to obtain images of 256x256 pixels. The MRI images obtained were analyzed with the ImageJ software.

Images were acquired Pre-gadolinium injection and 10-, 25-, 45-, 65-, 90- and 115-minutes Post-gadolinium injection. In vivo Pre- and Post- images were analyzed using ImageJ software (National Institutes of Health, Bethesda, Maryland). The intensity signal was measured in regions of interest, tumor region ($I_t$), contralateral region ($I_c$) and an area outside the mouse ($I_{noise}$). The Signal-to-noise ratio (SNR) of the region of interest were measured for each slice and time point and it is defined by $SNR_{(t)}=I_t/I_{noise}$, for the tumor region, and $SNR_{(c)}=I_c/I_{noise}$ for the contralateral region. The normalized enhancement ratio (NER%) of the tumor region to the contralateral region was calculated as $\%NER_t = \left[\left\{(SNR_t/SNR_c)_{post} - (SNR_t/SNR_c)_{pre}\right\}/(SNR_t/SNR_c)_{pre}\right] \times 100.$

*Determination of BBB integrity with Evans Blue extravasation*

For Evans blue infusion, 2ml/kg of Evans blue (E-2129, Sigma-Aldrich, MO) (2% dilution in PBS) was injected intravenously into the tail vein. After 30 min, the mice were deeply anesthetized with isoflurane and transcardially perfused in the heart beating through the left ventricle with 50 ml of ice-cold PB 0.1M followed by 50 ml of ice-cold 4% paraformaldehyde (PFA) in PB 0.1M. Brains were dissected afterwards.

*Chicken chorioallantoic membrane (CAM) assay*

For in vivo evaluation of the angiogenic inductive potential of SVZ derived CM, a CAM assay was performed as previously reported (Casas et al., 2018). Briefly, fertilized chicken eggs (Agricola Chorombo, Chile) were incubated at 38.5 °C with constant humidity. At embryonic day 1 (E1), 3 mL of albumin was extracted from each egg; a round window (2 cm2) was created on E4. A Bio-cellulose scaffold of 6mm of diameter was filled with 100 µl of medium to be assed: CM from SVZ-EGFRwt/amp or SVZ-EGFRvII, culture media (as negative control) and 100 µg VEGFA (positive control). On E8, the CAM vasculature was photographed; subsequently, each experimental condition scaffold was placed on top of the CAM; for each condition 14 eggs from different batches were used. On day E12, white cream was injected under the CAM before photographing every egg, in order to improve the visualization of the vessels.

*Inmunofluorescent (IF) and Inmunohistochemical (IHC) staining*

For immunostaining analyses, intracraneal tumors as well as human samples were fixed with 4% PFA for 12h at 4ªC and then tumors were embedded in paraffin. Paraffin sections (5 μM) were obtained with a microtome. Some animals were injected intraperitoneally with BrdU (Sigma Aldrich) (50mg/Kg) in saline solution 2 h before being sacrificed. Paraffin sections were incubated with primary antibodies (Supplementary Table S2) O/N at 4ºC. To detect BrdU, the paraffin sections were pre-incubated with heated 2N HCl for 15 min following by incubation in 0.1 M sodium borate [pH 8.5] during 10 min. The second day, sections were incubated with the appropriate secondary antibody. For IF, fluorescent antibodies (1:200 dilution) were used for 2h at room temperature. Prior to coverslip application, nuclei were counterstained with DAPI and imaging was done with Leica SP-5 confocal microscope. For IHC, sections were incubated with HRP conjugated antibodies (1:200 dilution) (Supplementary Table S2). Target proteins were detected with the ABC Kit and the DAB kit (Vector Laboratories).

*Western Blot analysis*

Protein extracts were prepared by re-suspending cell pellets or tumor tissue samples in lysis buffer (50 mM Tris (pH 7.5), 300 mM NaCl, 0.5% SDS, and 1% Triton X-100) and incubating the cells for 15 min at 100º. The lysed extracts were centrifuged at 13,000 g for 10 min at room temperature and the protein concentration was determined using a commercially available colorimetric assay (BCA Protein Assay Kit). Approximately 20 to 30 µg of protein were resolved by 10% or 12% SDS-PAGE and they were then transferred to a nitrocellulose membrane (Hybond-ECL, Amersham Biosciences). The membranes were blocked for 1 h at room temperature in TBS-T (10 mM Tris-HCl [pH 7.5], 100 mM NaCl, and 0.1% Tween-20) with 5% skimmed milk, and then incubated overnight at 4ºC with the corresponding primary antibody (Supplementary Table S2) diluted in TBS-T. Then, the membranes were incubated for 2 h at room temperature with their corresponding secondary antibody (HRP-conjugated anti mouse or anti rabbit, DAKO) diluted in TBS-T. Proteins were visible by enhanced chemiluminiscence with ECL (Pierce) using Amersham imager 680 and the signal was quantified by Fiji-ImageJ software.

*Quantitative reverse-transcriptase PCR (qRT-PCR)*

RNA was extracted from the tissue or the cell pellets using RNA isolation Kit (Roche) and it was digested with DNase I (Roche) according to the manufacturer's instructions. cDNA was synthesized with SuperScript II Reverse Transcriptase (Takara). qRT-PCR reactions were performed using the Light Cycler 1.5 (Roche) with the SYBR Premix Ex Taq (Takara). The primers used for each reaction are indicated in Supplementary Table S3. Gene expression was quantified by the double delta Ct method.

**Table S1. List of GBM cell lines.** The table indicates the genetic status of *EGFR* of the different primary GBM cell lines used in this study. (nd: not determined. 0: not present. 1: present)

| Cell line | Origin | EGFR amp | EGFR mut |
|---|---|---|---|
| **GBM1** | Hospital 12 de Octubre | 1 | 0 |
| **GBM2** | Hospital 12 de Octubre | 0 | 0 |
| **GBM3** | Hospital 12 de Octubre | 1 | 1 (EGFR vIII) |
| **GBM4** | Hospital 12 de Octubre | 1 | 1 (EGFR vIII) |
| **GBM5** | Hospital 12 de Octubre | 1 | 1 (EGFR vIII) |
| **GBM6** | Hospital 12 de Octubre | 1 | 1 (EGFR vIII) |
| **GBM7** | Hospital 12 de Octubre | 0 | 1 (V774M) |
| **GBM8** | Hospital 12 de Octubre | 0 | 1 |
| **GBM9** | Hospital 12 de Octubre | nd | nd |
| **GBM10** | Hospital 12 de Octubre | nd | nd |
| **GBM11** | Hospital 12 de Octubre | nd | nd |
| **GBM12** | Hospital 12 de Octubre | nd | nd |
| **GBM13** | Hospital 12 de Octubre | nd | nd |
| **GBM14** | Hospital 12 de Octubre | 0 | 0 |
| **GBM15** | Hospital 12 de Octubre | nd | nd |
| **GBM16** | Hospital 12 de Octubre | 1 | 0 |

**Table S2.** Antibodies

| Antibody | Dilution | Source |
|---|---|---|
| **α-SMA** | 1:500 (WB), 1:100 (IHC) | Santa Cruz Biotechnology |
| **β-Actin** | 1:1000 (WB) | Sigma |
| **β-catenin** | 1:1000 (WB) | Cell Signaling |
| **BrdU** | 1:100 | Dako |
| **AKT** | 1:1000 (WB) | Cell Signaling |
| **CD248** | 1:500 (WB), 1:100 (IF) | Santa Cruz Biotechnology, BDBiosciences |
| **CD34** | 1:100 (IF) | Leica |
| **EGFR** | 1:1000 (WB) | Cell Signaling |
| **EGFR vIII** | 1:1000 (WB) | Cell Signaling |
| **Endomucin** | 1:100 (IF) | Santa Cruz Biotechnology |
| **GAPDH** | 1:500 (WB) | Santa Cruz Biotechnology |
| **GFP** | 1:100(IF) | Santa Cruz Biotechnology |
| **HIF1α** | 1:500 (WB) | Santa Cruz Biotechnology |
| **PDGFRB** | 1:1000 (WB) | Cell Signaling |
| **p-AKT** | 1:1000 (WB) | Cell Signaling |
| **p-BMX** | 1:1000 (WB) | Invitrogen |
| **p-EGFR** | 1:1000 (WB) | Cell Signailing |
| **p-ERK** | 1:500 (WB) | Santa Cruz Biotechnology |
| **p-PDGFRB** | 1:1000 (WB) | Cell Signaling |
| **p-SRC** | 1:1000 (WB) | Cell Signaling |
| **p-STAT3** | 1:1000 (WB) | Cell Signaling |
| **SOX9** | 1:1000(WB) | Cell Signaling |
| **p-SOX9** | 1:1000 (WB) | Abcam |
| **Cy3 anti- mouse** | 1:200 (IF) | Jackson Immunoresearch |
| **Cy3 anti-rabbit** | 1:200 (IF) | Jackson Immunoresearch |

| | | |
|---|---|---|
| **Cy5 anti-mouse** | 1:200 (IF) | Jackson Immunoresearch |
| **Cy5 anti-rabbit** | 1:200 (IF) | Jackson Immunoresearch |
| **Cy5-anti rat** | 1:200 (IF) | Jackson Immunoresearch |
| **HRP anti-mouse** | 1:5000 (WB), 1:1000(IHC) | GE Healthcare |
| **HRP anti-rabbit** | 1:5000 (WB), 1:1000(IHC) | Santa Cruz Biotechnology |
| **HRP anti-rabbit** | 1:5000 (WB), 1:1000(IHC) | Cell Signaling |

**Table S3.** Primers used for the qRT-PCR analysis.

| Species | Gene | Forward (5'-3´) | Reverse (5'-3´) |
|---|---|---|---|
| **mouse** | **CD31** | TCCAGGTGTGCGAAATGCT | TGGCAGCTGATGCCTATGG |
| | **VE-CAD** | TTACTCAATCCACATACACATTTTCG | GCATGATGCTGTACTTGGTCATC |
| | **CD248** | TTGATGGCACCTGGACAGAGGA | TCCAGGTGCAATCTCTGAGGCT |
| | **αSMA** | ACCATCGGCAATGAGCGTTTCC | GCTGTTGTAGGTGGTCTCATGG |
| | **PDGFRB** | CCGGAACAAACACACCTTCT | TATCCATGTAGCCACCGTCA |
| | **MMP9** | GCAAGGGGCCGTGTCTGGAGATTC | GCCCACGTCGTCCACCTGGTT |
| | **LMNA** | TTGCCTCAACTGCAATGACAA | TCTCGATGTCGGTAAAACCCC |
| | **VEGFR1** | TTTGGCAAATACAACCCTTCAGA | GCAGAAGATACTGTCACCACC |
| | **VEGFR2** | CATCACCGAGAACAAGAACAAAACT | GATACCTAGCGCAAAGAGACACATT |
| | **TEK** | ACGGACCATGAAGATGCGTCAACA | TCACATCTCCGAACAATCAGCCTGG |
| | **EPHA2** | GCACAGGGAAAGGAAGTTGTT | CATGTAGATAGGCATGTCGTCC |
| | **AQP1** | AGGCTTCAATTACCCACTGGA | GTGAGCACCGCTGATGTGA |
| | **VEGF A** | TGCCAAGTGGTCCCAGGCTGC | CCTGCACAGCGCATCAGCGG |
| | **PDGF A** | GATACCTCGCCCATGTTCTG | CAGGCTGGTGTCCAAAGAAT |
| | **PDGF B** | GGGCCCGGAGTCGGCATGAA | AGCTCAGCCCCATCTTCATCTTACGG |
| | **PGF** | GAGGCCAGAAAGTCAGGGGGC | ATGGGCCGACAGTAGCTGCGA |
| | **NG2** | GACGGCGCACACACTTCTC | TGTTGTGATGGGCTTGTCAT |
| | **End** | TGCACTTGGCCTACGACTC | TGGAGGTAAGGGATGGTAGCA |
| | **Pi3kca** | GCTCTTCGCCATCACACAAAC | GGCATTCCTGTCATCAGCATC |
| | **Cxcr4** | GACTGGCATAGTCGGCAATGGA | CAAAGAGGAGGTCAGCCACTGA |
| | **Cx45** | ACTCCCTCTGTGATGTACCTGG | GTGCTGTTTCCAACGCATGGCA |
| | **Timp1** | TCTTGGTTCCCTGGCGTACTCT | GTGAGTGTCACTCTCCAGTTTGC |
| | **Chi3l1** | GCTTTGCCAACATCAGCAGCGA | AGGAGGGTCTTCAGGTTGGTGT |
| | **Lox** | CATCGGACTTCTTACCAAGCCG | GGCATCAAGCAGGTCATAGTGG |
| | **Serpine1** | CCTCTTCCACAAGTCTGATGGC | GCAGTTCCACAACGTCATACTCG |
| | **Cav1** | CACACCAAGGAGATTGACCTGG | CCTTCCAGATGCCGTCGAAACT |
| | **Car9** | GGCGAACGATTGAGGCTTCCTT | GCTGGTGACAGCAAAGAGAAGG |
| | **Car12** | TGGCTCTGAACACACCGTGAGT | TTGTCACTGGCGGTGCTGAAGT |
| | **Atf3** | GAAGATGAGAGGAAAAGGAGGCG | GCTCAGCATTCACACTCTCCAG |
| | **Hmox1** | CACTCTGGAGATGACACCTGAG | GTGTTCCTCTGTCAGCATCACC |
| **human** | **αSMA** | TAGCACCCAGCACCATGAAGATCA | GAAGCATTTGCGGTGGACAATGGA |
| | **NG2** | AGCTCTACTCTGGACGCC | ATCGACTGACAACGTGGC |
| | **CD248** | AGACCACCACTCATTTGCCTGGAA | AGTTGGGATAATGGGAAGCGTGGT |
| | **PDGFRB** | ACGGCTCTACATCTTTGTGCCAGA | TCGGCATGGAATGGTGATCTCAGT |
| | **SERPINE** | CATAGTGGAAGTGATAGAT | ACTCTGTTAATTCGTCTT |
| | **CD34** | CCTCAGTGTCTACTGCTGGTCT | GGAATAGCTCTGGTGGCTTGCA |
| | **VEGFA** | CTAACACTCAGCTCTGCCC | ACACACAAATACAAGTTGCCAA |
| | **ANGPT2** | ATTCAGCGACGTGAGGATGGCA | GCACATAGCGTTGCTGATTAGTC |
| | **IGFBP2** | CGAGGGCACTTGTGAGAAGCG | TGTTCATGGTGCTGTCCACGTG |
| | **CHI3L1** | CCACAGTCCATAGAATCCTCGG | TGCCTGTCCTTCAGGTACTGCA |
| | **LOX** | GATACGGCACTGGCTACTTCCA | GCCAGACAGTTTTCCTCCGCC |
| | **CAV1** | CCAAGGAGATCGACCTGGTCAA | GCCGTCAAAACTGTGTGTCCCT |
| | **CAIX** | GTGCCTATGAGCAGTTGCTGTC | AAGTAGCGGCTGAAGTCAGAGG |
| | **CAXII** | GACCTTTATCCTGACGCCAGCA | CATAGGACGGATTGAAGGAGCC |
| | **ATF3** | CGCTGGAATCAGTCACTGTCAG | CTTGTTTCGGCACTTTGCAGCTG |
| | **HMOX** | CCAGGCAGAGAATGCTGAGTTC | AAGACTGGGCTCTCCTTGTTGC |
| | **CD31** | AAGTGGAGTCCAGCCGCATATC | ATGGAGCAGGACAGGTTCAGTC |
| | **END** | GCAAGCACTTCAGCAACCAGCC | GGATCTGCCTTCCAGCACATTC |
| | **SOX9** | AGGAAGCTCGCGGACCAGTAC | GGTGGTCCTTCTTGTGCTGCAC |

**Supplementary Figures**

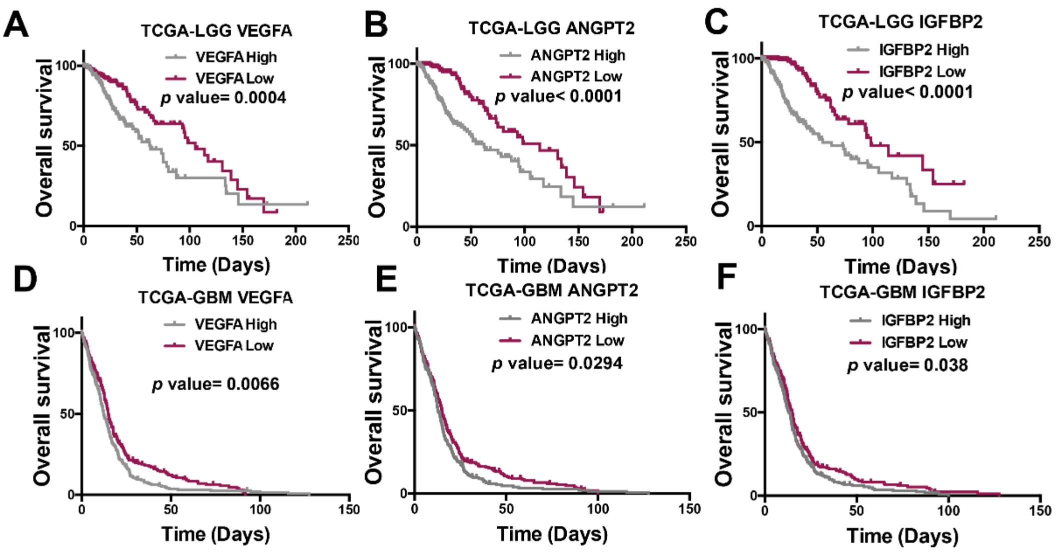


**Figure S1. Association of angiogenic molecules levels with glioma aggressiveness. A-C.** Kaplan-Meier overall survival curves of patients from the TCGA-LGG cohort (n=507). Patients were stratified into 2 groups based on high and low *VEGFA* (A), *ANGPT2* (B) or *IGFBP2* (C) expression values. **D-F.** Kaplan-Meier overall survival curves of patients from the TCGA-GBM cohort (n=525). Patients were stratified into 2 groups based on high and low *VEGFA* (D), *ANGPT2* (E) or *IGFBP2* (F) expression values.

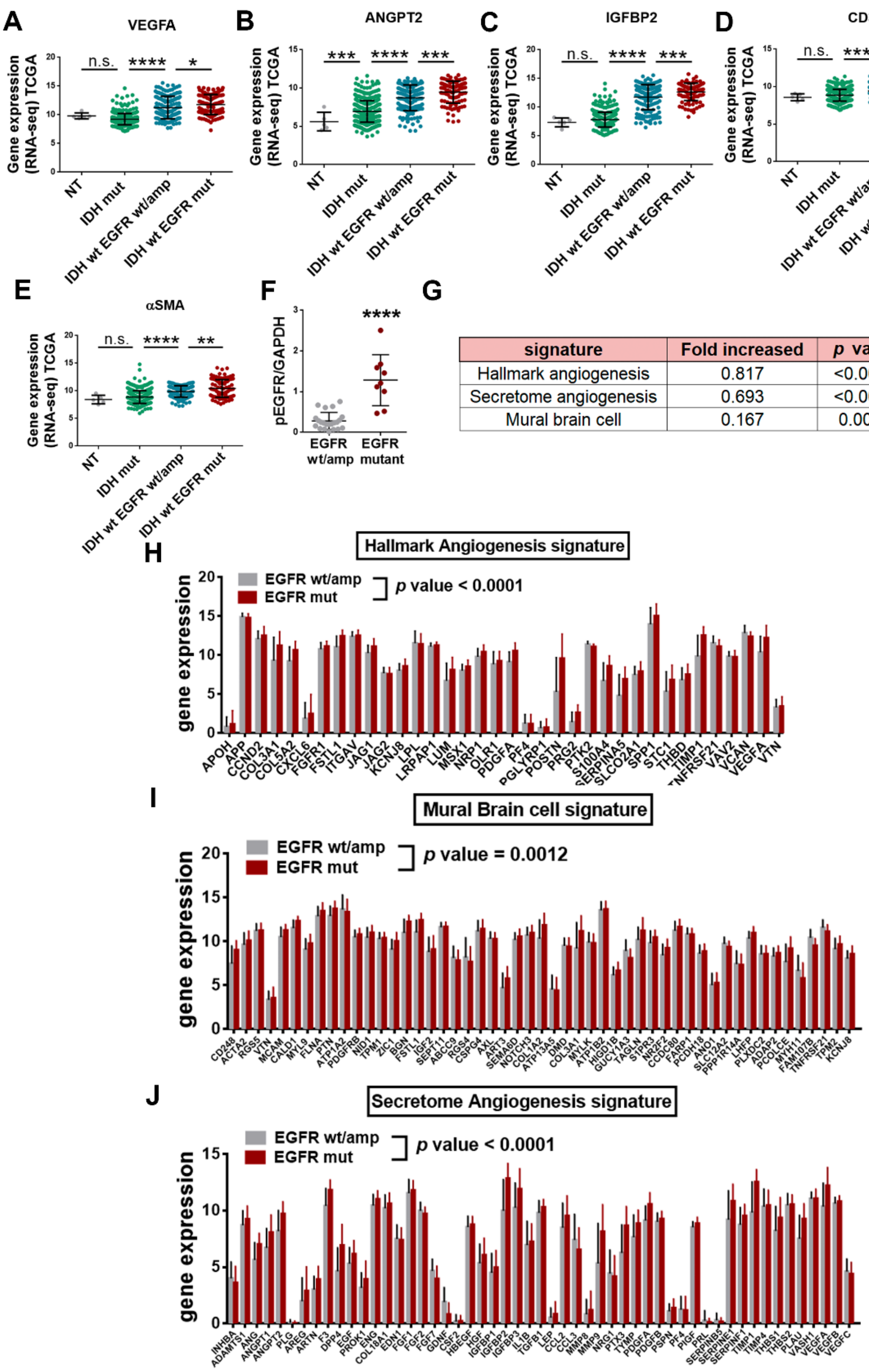


| signature | Fold increased | *p* value |
|---|---|---|
| Hallmark angiogenesis | 0.817 | <0.0001 |
| Secretome angiogenesis | 0.693 | <0.0001 |
| Mural brain cell | 0.167 | 0.0012 |

**Figure S2. Analysis of *EGFR-IDH* alterations and vascular molecules in human samples. A-E.** Analysis of mRNA levels of *VEGFA* (A), *ANGPT2* (B), *IGFBP2* (C), *CD34* (D) and *αSMA* (E) (RNAseq) in gliomas from the TCGA (LGG+GBM) cohort (n= 661). Tumors were stratified in three groups: IDHmut, IDHwt/EGFRwt/amp and IDHwt/GFRmut. **F.** Quantification of the WB analysis of pEGFR in patients stratified into 2 groups based on *EGFR* alterations (n=33). **G.** Summary of the expression of genes from each signature showed below (H-J). The table depicts the fold increase expression in EGFRmut compared to EGFRwt/amp tumors and the p value of the increment in the expression. **H-J.** Levels of expression of different signatures: hallmark angiogenesis (H), mural brain cell (I) and angiogenic secretome (J), in wt/amp or mut EGFR gliomas from the TCGA (LGG+GBM) cohort (n=319). *P ≤ 0.05; **P ≤ 0.01; ***P ≤ 0.001 ****P ≤ 0.0001; n.s. not significant.

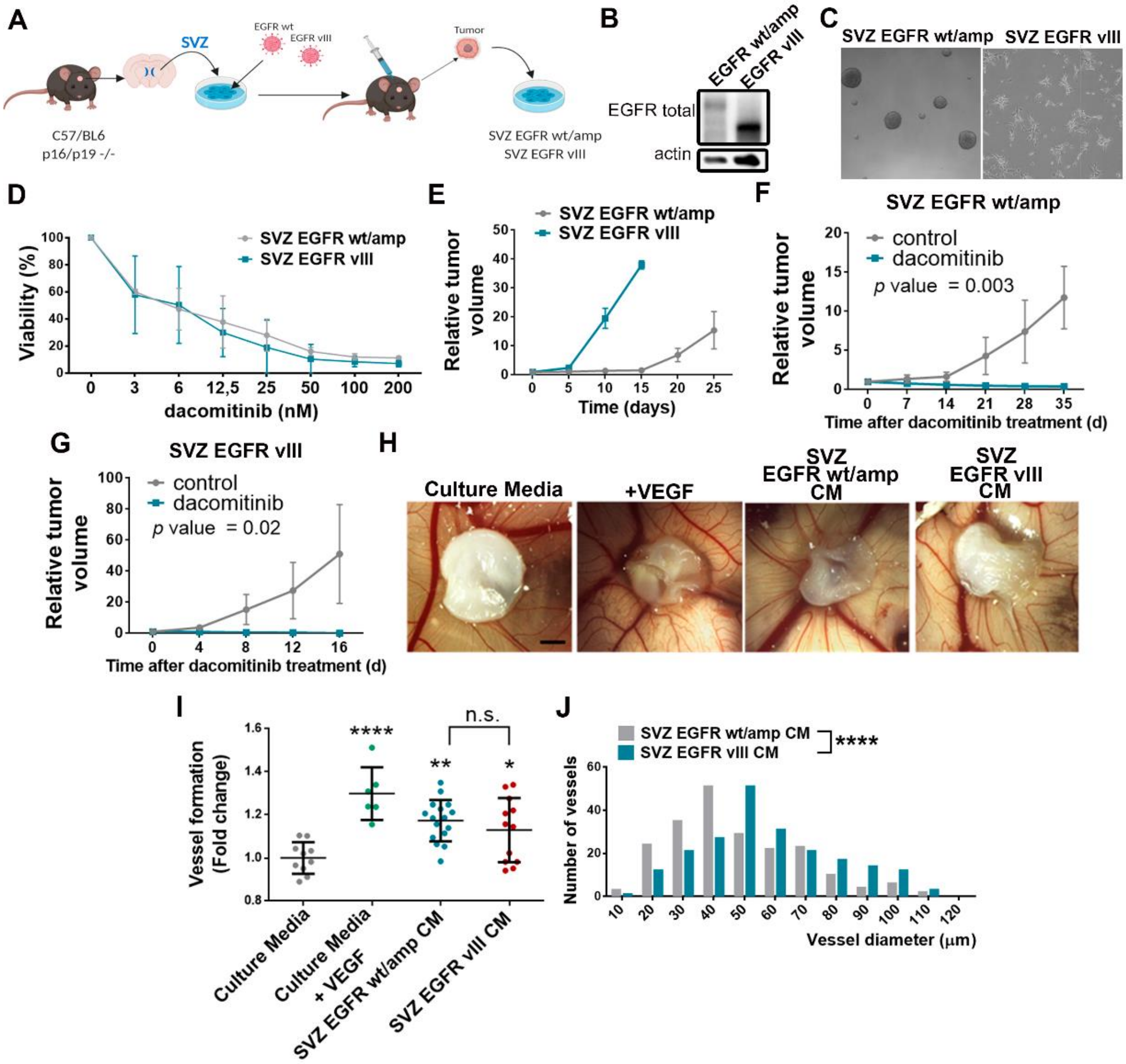


**Figure S3. Characterization of the murine glioma models: SVZ EGFR wt/amp and SVZ EGFR vIII. A.** Representative diagram of the generation of the SVZ-EGFRwt/amp and the SVZ-EGFRvIII murine glioma models. **B.** WB analysis of total EGFR in SVZ cell lines. Actin expression was used for normalization. **C.** Representative images of SVZ cell lines grown *in vitro*. **D.** Viability (represented in % related to the control) of SVZ cell lines grown *in vitro* in the presence of different concentrations of dacomitinib. **E.** SVZ-EGFRwt/amp and SVZ-EGFRvIII were implanted subcutaneously in the nude mice and tumor growth was measured with a caliper. The graph represents the fold increase in tumor volume (n=3). **F-G.** SVZ-EGFRwt/amp (F) and SVZ-EGFRvIII (G) cells were implanted subcutaneously in the nude mice. When tumors became visible, animals were treated with dacomitinib (15mg/Kg/day). The graphs represent the fold increase in tumor volume (n=5). **H.** Representative images of blood vessel formation around the bio-cellulose scaffolds. Scaffolds were incubated with culture media, culture media with VEGF (as a positive control) and culture media plus conditioned media (CM) from SVZ-EGFRwt/amp or SVZ-EGFRvIII cells. **I-J.** Analysis of the number of vessels (I) and the vessel diameter (J) in each condition in (H). ****P ≤ 0.0001, n.s. not significant. Scale bar: 100 μm.

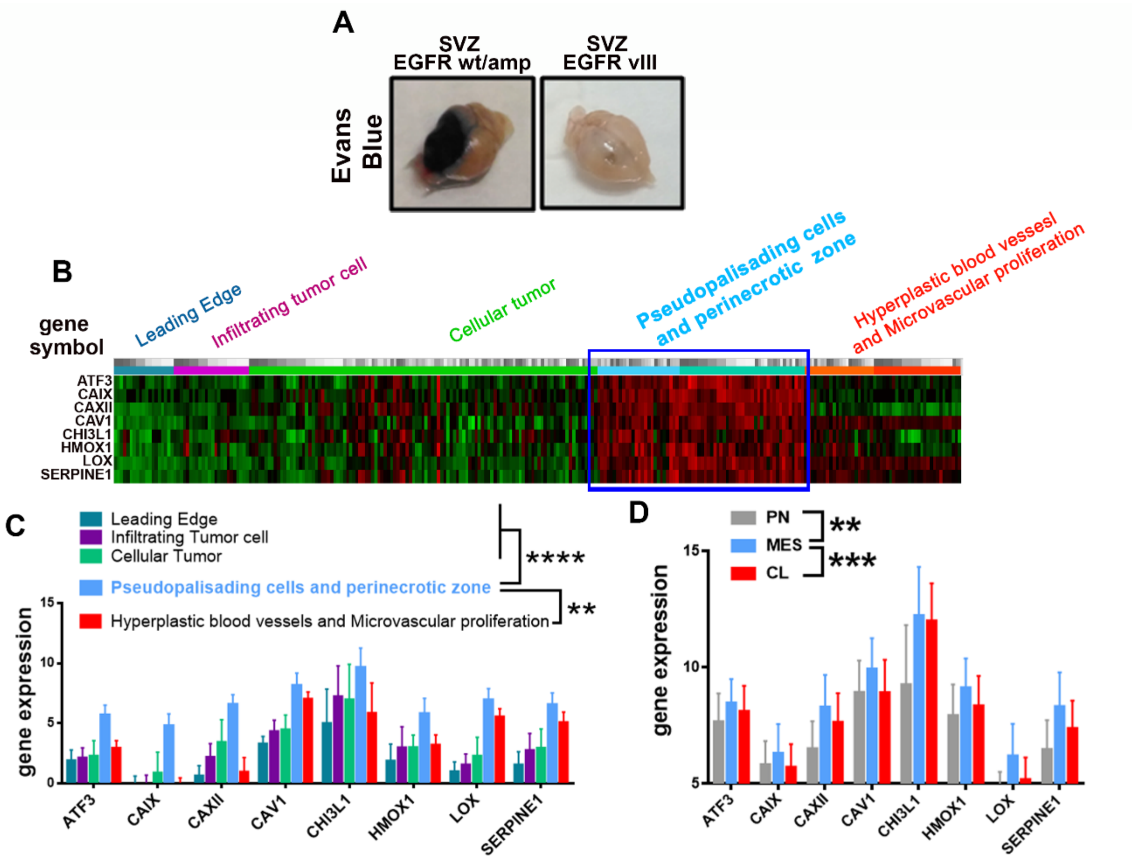


**Figure S4. Association of EGFR alterations with BBB leakage and definition of a hypoxia signature in gliomas. A.** Representative image of the whole brain of mice harboring SVZ-EGFRwt/amp or SVZ-EGFRvIII gliomas tumor showing Evans Blue extravasation. **B.** Selection of the most relevant genes of the hypoxia and the HIF1α pathway signatures that were up-regulated in the perinecreotic and pseudopalisading cell necrosis zones. YvyGap (IVY Glioblastoma atlas project) data set analysis was used**. C.** Levels of expression of the hypoxic-related signature in gliomas of the TCGA (GBM) cohorts depending on the different anatomical regions of the tumors. **D.** Levels of expression of the hypoxic-related signature in gliomas of the TCGA (GBM) cohort according to the molecular subtype's classification. PN: proneural, MES: mesenchymal, CL: classic. **P ≤ 0.01; ***P ≤ 0.001: ****P ≤ 0.0001

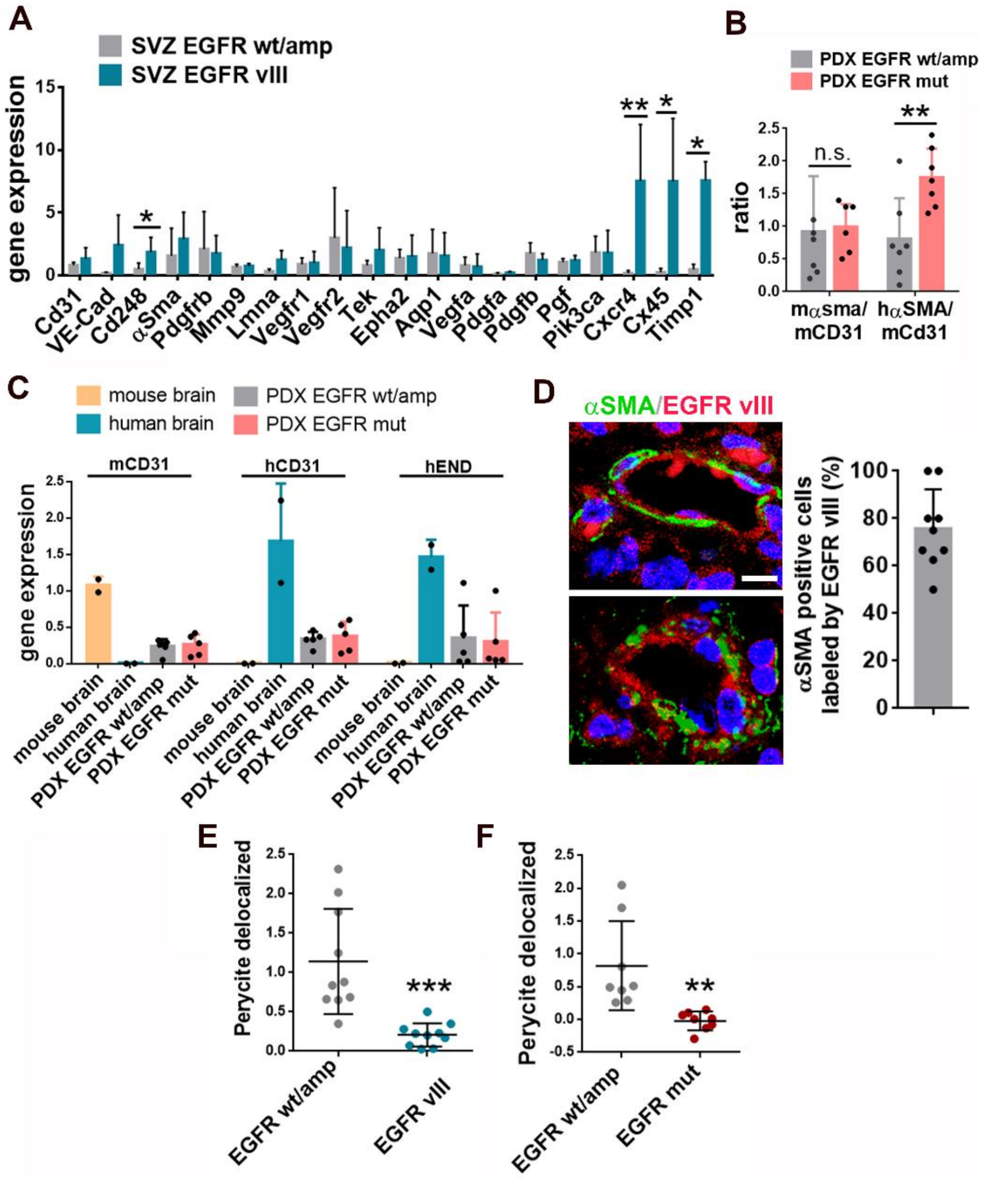


**Figure S5. Participation of EGFR mutations in the formation of glioma-derived pericytes. A.** qRT-PCR analysis of angiogenesis-related genes in SVZ-EGFRwt/amp and SVZ-EGFRvIII cells. *Actin* was used for normalization (n=3). **B.** Ratio of the expression of pericyte (mouse and human) to endothelial (mouse) genes (qRT-PCR analysis) in PDXs models expressing wt/amp or mut EGFR (n=7). **C.** qRT-PCR analysis of endothelial–related genes in EGFRwt/amp and EGFRmut tumor xenografts. Human or mouse tissue was used as control. *HPRT* or A*ctin* was used for normalization (n=5). **D.** Representative images of αSMA and EGFRvIII immunofluorescent (IF) staining of sections from a tumor sample expressing EGFRvIII. The graph on the right shows the quantification of the fraction of pericytes carrying the EGFRvIII genetic alteration per field (n=9). **E-F.** Quantification of the number of delocalized pericytes delocated in SVZ (E) or PDX (F) tumor sections. *P ≤ 0.05; **P ≤ 0.01; ***P ≤ 0.001; n.s. not significant. Scale bar: 10 μm.

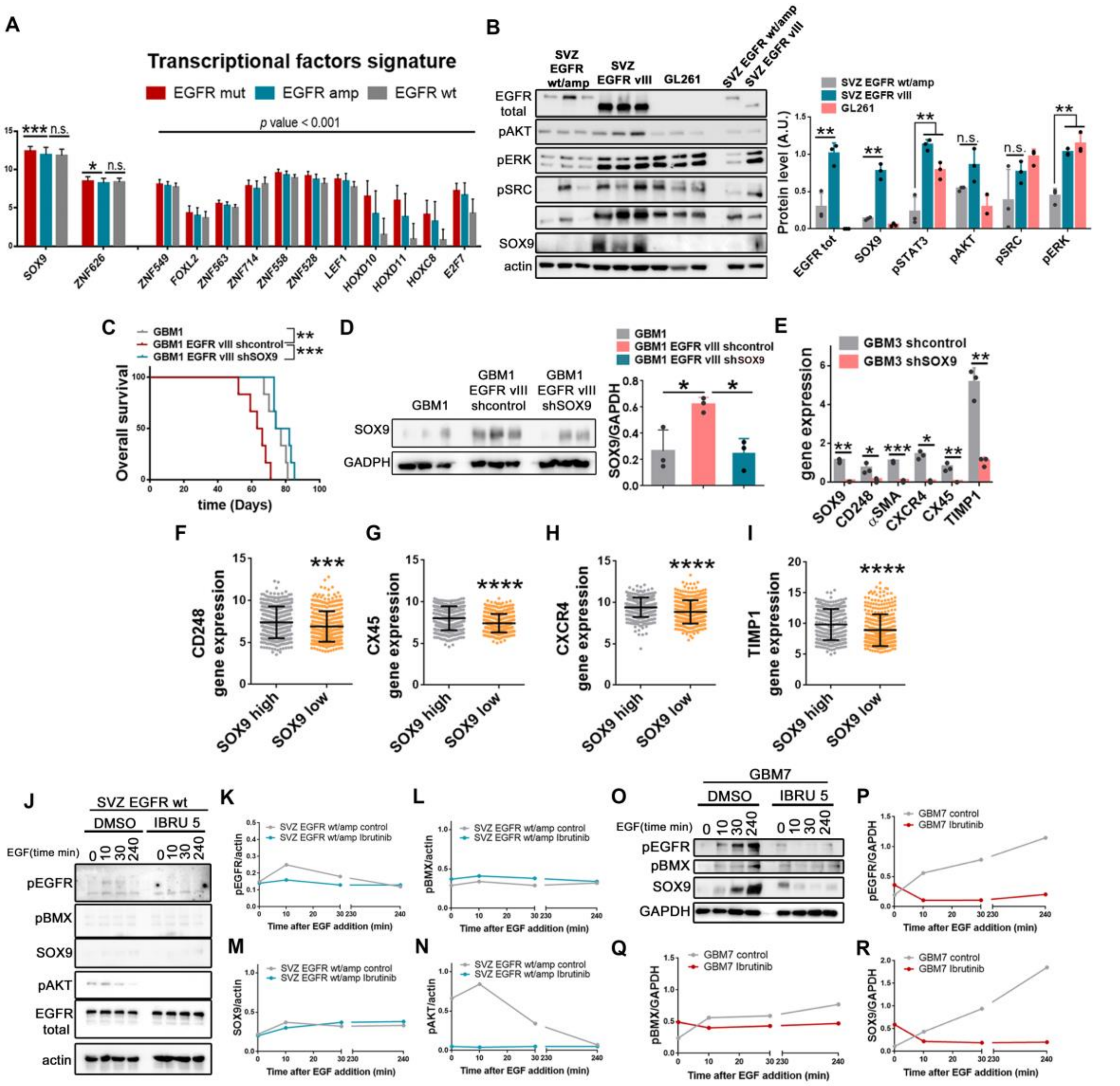


**Figure S6. Correlation of SOX9 expression with EGFR mutations in gliomas. A.** Analysis of the expression of different transcription factors (RNAseq) in gliomas from the TCGA GBM+LGG cohort, n=306, grouped according to the genetic status of *EGFR*. **B.** WB analysis and quantification of EGFR and SOX9 in tumor tissue extracts from SVZ and GL261 tumors. Extracts from SVZ cells were used as controls and loaded on the right. The expression of Actin was used for normalization. **C.** Kaplan-Meier overall survival curves of mice that were orthotopically injected with GBM1, GBM1-EGFRvIII and GBM1-EGFRvIII-shSOX9 (n=6). **D.** WB analysis of SOX9 expression in tumor tissue extracts from (C). GAPDH expression was used for normalization (n=3). **E.** qRT-PCR analysis of *SOX9*, *CD248*, *αSMA*, *CXCR4*, *CX45* and *TIMP1* in the tumors formed by GBM3-shControl or GBM3-shSOX9 cells. The expression of *HPRT* was used for normalization (n=3). **F-I.** Analysis of *CD248* (F), *CX45* (G), *CXCR4* (H) and *TIMP1* (I) mRNA expression (RNAseq) in gliomas from the TCGA LGG+GBM cohort (n=702). Tumors were classified in two groups based on high or low SOX9 expression values. **J.** WB analysis and quantification of pEGFR, pBMX, SOX9, pAKT and total EGFR total in extracts from SVZ-EGFRwt/amp cells incubated with EGF (100ng/ml) for the times indicated, in the presence of DMSO or Ibrutinib (5µm). Actin was used as loading control. **K.** WB analysis and quantification of pEGFR, pBMX and SOX9 in extracts from GBM7 cells incubated with EGF (100ng/ml) for the times indicated, in the presence of DMSO or ibrutinib (5µM). GAPDH was used as loading control. *P ≤ 0.05; **P ≤ 0.01; ***P ≤ 0.001;****P ≤ 0.0001; n.s. not significant.
